# The *NORAD*-pumilio regulatory axis links lncRNA dysregulation to tau propagation-associated phenotypes

**DOI:** 10.64898/2026.06.30.735697

**Authors:** Joseph E. Zemke, Guangming Huang, Emma Starr, Matthew Broder, Jacob Marsh, Arun Renganathan, Bridget Phillips, Thomas W. Marsh, Miguel Minaya, Carlos Cruchaga, Abhirami K. Iyer, Dominantly Inherited Alzheimer Network, Celeste M. Karch

## Abstract

Long non-coding RNAs (lncRNAs) are increasingly implicated in neurodegenerative disease, yet their roles in tauopathy remain poorly understood. Here, we defined the lncRNA landscape across iPSC-derived neurons, astrocytes, and microglia harboring the frontotemporal dementia-associated *MAPT* IVS10+16 mutation and investigated how lncRNA dysregulation interfaces with tau pathology. Transcriptomic analyses revealed extensive cell-type specific lncRNA expression changes, with neurons exhibiting the greatest degree of mutation-associated remodeling. Comparative analyses with *MAPT* IVS10+16 patient brain tissue identified *NORAD* and *MIR22HG* as lncRNAs significantly dysregulated across all three cell types and human brains. *NORAD* was also altered in Alzheimer’s disease and Parkinson’s disease brains, suggesting a broader role in neurodegenerative disease. Mechanistically, *NORAD*-associated protein networks converged on pathways related to RNA regulation, cytoskeletal organization, proteostasis, and tau interaction networks. Given the established role of *NORAD* in regulating PUM1 and PUM2 RNA-binding (pumilio) proteins, we examined the *NORAD*-pumilio axis and identified enrichment of pumilio-associated pathways linked to autophagy, endocytosis, proteostasis, and cytoskeletal regulation. *NORAD* depletion reduced tau seeding and uptake, whereas functional depletion of *PUM1* or *PUM2* increased both processes, supporting an antagonistic relationship between *NORAD* and pumilio signaling in modulation of tau aggregation. Together, these findings identify widespread lncRNA dysregulation across neural cell types in the setting of a *MAPT* mutation and nominate the *NORAD*-pumilio axis as a regulatory pathway linking RNA homeostasis and tau propagation biology.

## Introduction

Tauopathies are a heterogeneous group of neurodegenerative disorders characterized by the abnormal accumulation and propagation of tau aggregates throughout the brain^1^. Frontotemporal lobar degeneration with tau pathology (FTLD-tau) is a primary tauopathy and can be caused by dominantly inherited mutations in the *microtubule-associated protein tau* (*MAPT*) gene^2, 3^. The FTLD-tau causing *MAPT* IVS10+16 mutation alters *MAPT* splicing and promotes increased production of 4R tau isoforms, ultimately leading to progressive tau aggregation and neurodegeneration^3–7^. Increasing evidence suggests that tau pathology propagates through interconnected neural networks via cellular uptake and intracellular seeding of pathogenic tau species, processes believed to contribute to disease progression^8–11^. While substantial progress has been made in defining the biochemical mechanisms underlying tau aggregation^12, 13^, the upstream regulatory pathways controlling cellular susceptibility to tau uptake and seeding remain incompletely understood.

Long non-coding RNAs (lncRNAs) are emerging as important regulators of cellular identity, RNA metabolism, and stress-response signaling within the nervous system^14, 15^. Although lncRNAs do not encode proteins, they regulate diverse biological processes through interactions with chromatin, RNA-binding proteins, mRNA transcripts, and signaling pathways^16–19^. Notably, lncRNAs often exhibit substantially greater cell-type specificity than protein-coding genes and are highly enriched within the brain^20^. Dysregulation of lncRNAs has been implicated across multiple neurodegenerative disorders, including Alzheimer’s disease (AD), Parkinson’s disease (PD), and amyotrophic lateral sclerosis (ALS)^21–26^. Recent studies have begun to implicate lncRNAs in tauopathy biology^27–29^. Prior work from our group demonstrated that *MAPT* mutations broadly alter lncRNA expression in human iPSC-derived neurons and identified lncRNAs that regulate stress granule biology, autophagy, and tau aggregation-associated pathways^21, 22^. Other groups have similarly implicated lncRNAs, including *NEAT1*, *MALAT1*, *MAPT-AS1*, and *BACE1-AS* in RNA metabolism, proteostasis, inflammatory signaling, and tau-related neurodegenerative processes^23, 24, 30, 31^. Together, these studies support the concept that non-coding RNA biology represents an important and underexplored layer of neurodegenerative disease pathobiology. However, how lncRNAs influence tau propagation-associated phenotypes across distinct neural cell types remains poorly understood.

Increasing evidence suggests that tau propagation and neurodegeneration are influenced not only by tau aggregation itself, but also by broader cellular pathways regulating RNA homeostasis, proteostasis, vesicular trafficking, and cytoskeletal organization^32–35^. RNA-binding proteins and post-transcriptional regulatory networks have emerged as important modulators of neuronal vulnerability across neurodegenerative diseases, including tauopathies, synucleinopathies, and ALS^36–41^. In parallel, pathways governing autophagy, endocytosis, stress granule dynamics, and actin cytoskeleton remodeling have been increasingly implicated in tau uptake, intracellular seeding, and aggregate propagation^21, 35, 42–44^. Notably, many lncRNAs function through interactions with RNA-binding proteins and regulatory protein complexes^19, 38, 45^, positioning them as potential upstream coordinators of multiple disease-relevant pathways. However, whether lncRNA dysregulation contributes to the intersection of RNA regulatory, cytoskeletal, and tau propagation-associated pathways across neural cell types remains largely unknown.

Importantly, these disease-associated pathways operate across multiple neural cell types that differentially contribute to tauopathy progression. Tauopathies involve complex interactions among neurons, astrocytes, and microglia, each contributing differently to tau aggregation, propagation, and neuroinflammatory responses^46, 47^. Neurons represent the primary site of tau accumulation^48^, while astrocytes and microglia influence extracellular tau clearance, inflammatory signaling, and propagation-associated pathways^49–52^. Importantly, lncRNA expression is highly cell-type specific, suggesting that distinct lncRNA regulatory programs may differentially shape disease-associated processes across neural cell populations^20^. Despite this, most studies examining lncRNAs in neurodegeneration have focused predominantly on neurons, leaving the contribution of astrocytic and microglial lncRNA biology comparatively unexplored.

Here, we defined the lncRNA landscape across iPSC-derived neurons, astrocytes, and microglia harboring the FTLD-associated *MAPT* IVS10+16 mutation and integrated these findings with human brain transcriptomic datasets. We identified widespread cell-type specific lncRNA dysregulation and nominate *NORAD* as a disease-associated lncRNA altered across multiple neurodegenerative diseases. Mechanistically, we identify the *NORAD*–pumilio regulatory axis as a pathway linking RNA homeostasis, cytoskeletal regulation, and tau propagation-associated phenotypes, including tau uptake and seeding. Together, these findings position lncRNA-mediated regulatory networks as an important and underexplored dimension of tauopathy biology.

## Results

### LncRNA expression in neurons, astrocytes, microglia and brain homogenates

Considering the strong cell-type specificity of lncRNA biology^20, 53–55^, we first defined the baseline lncRNA expression landscape across iPSC-derived neurons, astrocytes, and microglia (**Figure 1A**). Re-analysis of previously generated ribo-depleted RNA-sequencing datasets from *MAPT* WT cells^21, 22, 56–58^, using stringent expression thresholds (≥0.1 TPM in ≥10% of samples), revealed a lncRNA architecture composed of both shared and highly cell-type restricted programs (**Figure 1B; Supplemental Table 1**). iPSC-derived microglia exhibited a markedly expanded repertoire of cell-type specific lncRNAs (n=3,269), compared to iPSC-derived neurons (n=1,175) and iPSC-derived astrocytes (n=560), consistent with the transcriptionally dynamic functions of microglia that may require more specialized lncRNA repertoire in this cell type. In parallel, 2,838 lncRNAs were conserved across all three cell types, defining a core lncRNA program shared across cell types that may play roles in fundamental cellular processes (**Figure 1B**).

**Figure 1:**
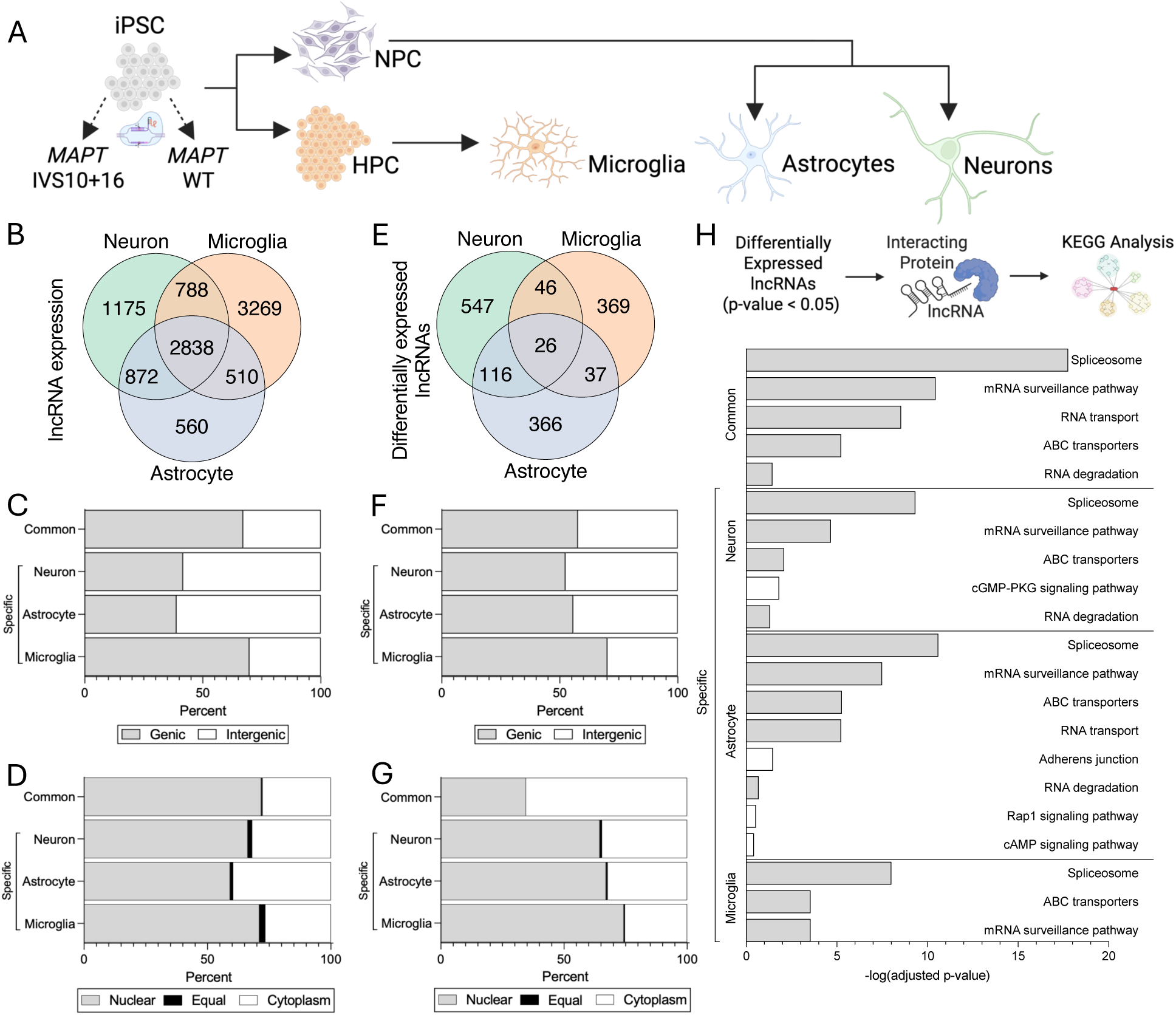
Cell type-resolved lncRNA profiling identifies shared and cell-specific programs altered by *MAPT* IVS10+16. A. Isogenic iPSCs were created by correcting patient derived *MAPT* IVS10+16/WT iPSCs to *MAPT* WT/WT using CRISPR/Cas9. Isogenic iPSCs were differentiated into neurons, astrocytes, and microglia. RNA-sequencing with ribodepletion was performed and protein coding RNA and lncRNA were quantified. B. Expression of lncRNAs in iPSC-derived neurons (n=5673), astrocytes (n=4780), and microglia (n=7405) were defined based on transcripts present in at least 10% of samples with expression of ≥ 0.1 TPM. A subset of lncRNAs were detected in all cell types (common; n=2838). C. Proportion of genic (gray) and intergenic (white) cell-type specific and commonly expressed lncRNAs. D. Subcellular localization of cell-type specific and commonly expressed lncRNAs. Nuclear (gray), cytoplasm (white), and equally distributed (black). E. Differentially expressed lncRNAs in *MAPT* IVS10+16 compared with isogenic controls (p≤0.05) were identified in iPSC-derived neurons (n= 735), astrocytes (n=545), and microglia (n=478). F. Proportion of genic (gray) and intergenic (white) cell-type specific and commonly dysregulated lncRNAs. G. Subcellular localization of cell-type specific and commonly dysregulated lncRNAs. Nuclear (gray), cytoplasm (white), and equally distributed (black). H. Pathway analysis of cell-type specific and commonly dysregulated lncRNA-protein interactions. lncRNA-protein interactions were determined using predicted and experimentally validated interactions (FDR≤0.05). KEGG analysis was conducted on those proteins identified. Unique KEGG pathway (white).

To begin to infer the functional roles of these lncRNA programs, we annotated transcripts based on genomic context and subcellular localization. While most lncRNAs remain incompletely characterized, genomic and subcellular localization provide general context for potential regulatory modes. For example, genic lncRNAs are often associated with *cis* regulation, while intergenic lncRNAs have *trans*-acting functions^59^. Across cell types, neurons and astrocytes were enriched for intergenic lncRNAs (**Figure 1C; Supplemental Table 2; Supplemental Table 3**; 58% and 61%, respectively), whereas microglia and the shared lncRNA pool were predominantly genic lncRNAs (**Figure 1C**; **Supplemental Table 4; Supplemental Table 5;** 69% and 67%, respectively), suggesting differences in genomic organization of lncRNAs across cell types. Notably, lncRNAs in all groups were strongly enriched in the nucleus (**Figure 1D; Supplemental Tables 6-9;** neuron-specific: 66%; astrocyte-specific: 59%; microglia-specific: 70%; common: 71%), based on lncATLAS annotations^60^. This nuclear localization is consistent with established roles for lncRNAs in transcriptional regulation, chromatin organization, and RNA processing and suggests a predominant engagement of nuclear regulatory mechanisms relative to cytoplasmic functions. Together, these analyses define a cell-type resolved and functionally informed lncRNA landscape and highlight microglia as a cell type with an expanded lncRNA repertoire. These data establish a framework to assess whether *MAPT* mutations preferentially disrupt lncRNA regulation in a cell-type-dependent manner.

### MAPT IVS10+16 disrupts lncRNA expression in neurons, astrocytes, and microglia

Having established a cell-type resolved lncRNA landscape, we next asked how the FTLD-tau-causing *MAPT* IVS10+16 mutation perturbs these programs. We re-analyzed RNA-sequencing datasets from isogenic iPSC-derived neurons, astrocytes, and microglia harboring either *MAPT* WT or *MAPT* IVS10+16 (**Figure 1A**). Differential expression analysis revealed both shared and cell-type specific lncRNA dysregulation across all three cell types (p≤0.05; **Figure 1E; Supplemental Table 10**). *MAPT* IVS10+16 expressing iPSC-derived neurons exhibited the greatest number of differentially expressed, cell-type specific lncRNAs (n=547), followed by astrocytes (n=366) and microglia (n=369; **Figure 1E**). The enrichment of detectable lncRNA changes in neurons in response to *MAPT* perturbation could reflect the primary vulnerability of neurons in tauopathies. Only 26 lncRNAs were commonly dysregulated across all three cell types (**Figure 1E**), highlighting cell-type restricted responses to *MAPT* perturbation. Among shared signatures, neurons and astrocytes displayed the greatest overlap (n=116), whereas overlap with microglia was more limited (**Figure 1E**), consistent with shared lineage relationships or regulatory programs between neurons and astrocytes.

We next annotated the dysregulated lncRNAs in each cell type to infer functional consequences using genomic context and subcellular localization. Cell-type specific differentially expressed lncRNAs were enriched for intergenic transcripts in neurons and astrocytes, whereas differentially expressed lncRNAs in microglia and commonly dysregulated lncRNAs were primarily genic (**Figure 1F; Supplemental Tables 11-14**), mirroring baseline organizational differences (**Figure 1C**). Most dysregulated lncRNAs localized to the nucleus (**Figure 1G**; **Supplemental Tables 15-17**; neurons: 64%; astrocytes: 67%; microglia: 74%), supporting a primary role in transcriptional and chromatin-associated regulation. Interestingly, the small set of commonly dysregulated lncRNAs (n=26) were enriched in the cytoplasm (**Figure 1G**; **Supplemental Table 18;** 65%), suggesting a shared disruption of post-transcriptional regulatory processes across cell types.

To further define the function of dysregulated lncRNAs, we mapped lncRNA-protein interactions using the lncSEA2.0 database and performed KEGG pathway enrichment on significantly associated protein interactors (**Figure 1H**; FDR≤0.05)^61^. Across cell types, proteins interacting with dysregulated lncRNAs converged on core pathways, including the spliceosome, mRNA surveillance, ABC transporters, and RNA transport and degradation (**Figure 1H**), indicating shared disruption of RNA processing and cellular homeostasis by *MAPT* IVS10+16. Cell-type specific signatures also emerged: neuronal lncRNAs interacted with proteins enriched in cGMP–PKG signaling, while astrocytic lncRNAs interacted with proteins associated with adherens junction, Rap1, and cAMP signaling pathways (**Figure 1H**). Many of these pathways are implicated in neurodegeneration^62–69^. Together, these data demonstrate that the *MAPT* IVS10+16 mutation induces both convergent and cell-type specific lncRNA expression changes, implicating RNA processing and signaling pathways associated with neurodegenerative biology.

### Integrated iPSC and human brain analyses prioritize NORAD as a disease-associated lncRNA

To assess the translational relevance of these findings, we next asked whether lncRNAs identified in our cell-based models are represented in human brain tissue. We compared both common and cell-type specific differentially expressed lncRNAs with those detected in human brains. Notably, 96% of commonly dysregulated lncRNAs were detected in human brains, along with high representation of neuron-specific (90%) and astrocyte-specific (85%) lncRNAs, whereas microglia-specific lncRNAs showed lower representation (68%; **Supplemental Table 19**). The lower representation among microglia likely reflects the limited capture of microglia in bulk brain RNA-sequencing accompanied by the lower abundance of lncRNA expression overall^70^. Together, our findings suggest that a large fraction of lncRNA changes identified *in vitro* are preserved in human brain.

Having identified *MAPT* IVS10+16-associated lncRNA changes across neurons, astrocytes, and microglia, we next asked which of these alterations are also present in human brain tissue harboring the same mutation. By comparing cell-type specific lncRNA signatures with those detected in *MAPT* IVS10+16 patient brain homogenates (p≤0.05), we identified substantial overlap across model systems, with neurons exhibiting the greatest concordance with human brain (n=55), followed by astrocytes (n=34) and microglia (n=27) (**Figure 2A**). To further characterize these shared candidates, we examined their expression patterns across iPSC-derived neurons, astrocytes, microglia, and *MAPT* IVS10+16 patient brains (**Figure 2B**). *CASC9* was significantly downregulated across cell types but unchanged in brain tissue, while *MIR4458HG, AC112178.1, LINC00884, AL137782.1, and LINC02693* were significantly upregulated in all cell types and unchanged in brain tissue (**Figure 2B**). Interestingly, we observed *AC093673.1*, *HCG11*, *KDM7A-DT*, *MIR4435-2HG*, *MIR503HG*, *SNHG16*, and *ZNF667-AS1* were significantly reduced in *MAPT* mutant neurons and significantly upregulated in mutant astrocytes and microglia (**Figure 2B**), which may suggest a cell-type specific role for these lncRNAs. However, these lncRNAs were not differentially expressed in brain homogenates from *MAPT* IVS10+16 carriers. Notably, among the 26 lncRNAs commonly altered across all three cell types, only *NORAD* and *MIR22HG* were also differentially expressed in *MAPT* IVS10+16 patient brain tissue (**Figure 2B**). *NORAD* and *MIR22HG* displayed opposing effects in neuronal and glial populations, with reduced expression in neurons and increased expression in astrocytes and microglia, suggesting cell-type dependent responses to a *MAPT* mutation. Expression of *NORAD* and *MIR22HG* in *MAPT* IVS10+16 patient brains was consistent with *MAPT* mutant neurons (**Figure 2B**). Together, these findings position *NORAD* and *MIR22HG* as high-confidence disease-associated lncRNAs.

**Figure 2:**
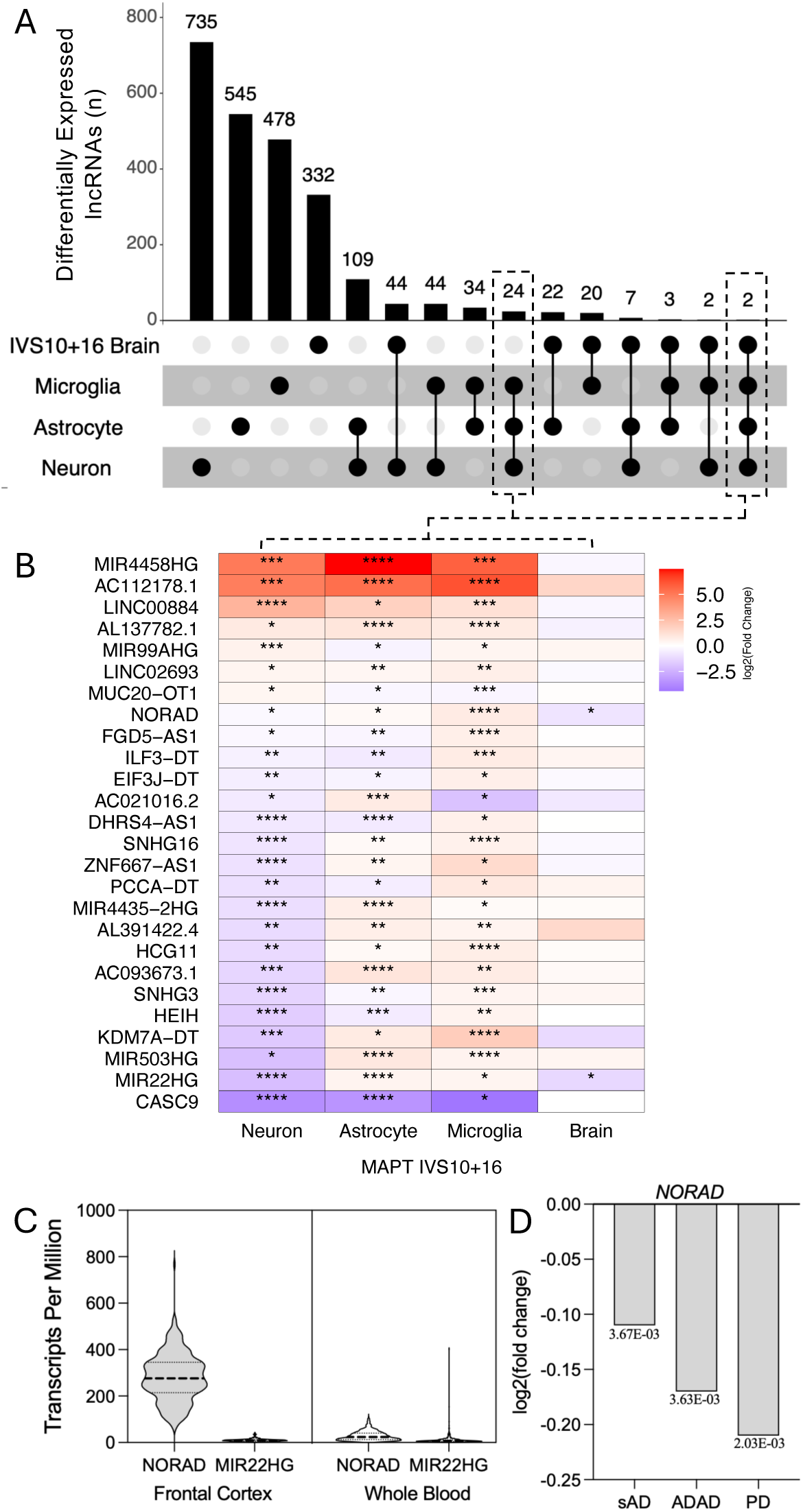
Human brain integration identifies *NORAD* as a brain-enriched lncRNA dysregulated across tauopathy and neurodegenerative disease. A. UpSet plot representing lncRNAs dysregulated as a function of the *MAPT* IVS10+16 mutation in iPSC-derived neurons, astrocytes, microglia, and the human brain tissues from *MAPT* IVS10+16 compared to controls (p≤0.05). B. Heatmap representing log_2_(fold-change) of 26 dysregulated lncRNAs shared in *MAPT* IVS10+16 neurons, astrocytes, microglia, and brains. *, p≤0.05; **, p<0.01; ***, p<0.001; ****, p<0.0001. C. *NORAD* and *MIR22HG* gene expression (transcripts per million) in the frontal cortex and whole blood (GTEX). D. *NORAD* is significantly downregulated in sporadic Alzheimer’s Disease (sAD) brains, autosomal dominant Alzheimer’s Disease (ADAD) brains, and Parkinson’s Disease (PD) brains compared to healthy controls.

Because *NORAD* and *MIR22HG* were the only lncRNAs differentially expressed across all three cell types and in *MAPT* IVS10+16 patient brains, we sought to further characterize their expression patterns and tissue specificity using GTEx (v11), a comprehensive resource of tissue- and cell-specific expression in humans. Despite these shared directional changes, the two lncRNAs exhibited markedly different expression profiles across human tissues. *NORAD* was highly enriched in brain regions vulnerable to FTLD, including the frontal cortex (**Figure 2C**; **Supplemental Figure 1A**), while showing low expression in the periphery (whole blood), consistent with a predominantly brain-associated expression pattern. In contrast, *MIR22HG* displayed comparatively low expression across brain regions and similarly low expression levels in brain and blood (**Figure 2C**; **Supplemental Figure 1B**).

To determine whether *NORAD* dysregulation extends beyond FTLD-tau, we first investigated secondary tauopathies and found that *NORAD* expression was significantly reduced in sporadic AD brains (p=3.67×10^−3^) and autosomal dominant AD brains (p=3.63×10^−3^) relative to controls (**Figure 2D**; **Supplemental Table 20**). Because PD exhibits widespread proteostatic dysfunction and can develop secondary tau pathology^71^, we next assessed *NORAD* expression in PD brains and similarly observed significantly reduced *NORAD* expression (p=2.03×10^−3^; **Figure 2D**; **Supplemental Table 20**). *MIR22HG* was only nominally significantly downregulated in the autosomal dominant AD brains (p=0.02; **Supplemental Table 20**). Together, these findings suggest that *NORAD* dysregulation extends across multiple neurodegenerative diseases characterized by proteostatic dysfunction and tau-associated pathology, prioritizing it for further mechanistic investigation.

### NORAD interaction networks converge with tau-associated cytoskeletal and proteostasis pathways

We next sought to better understand the molecular pathways potentially associated with *NORAD*’s function in neurodegeneration. LncRNAs frequently exert their functions through interactions with RNA-binding and regulatory proteins. Therefore, we leveraged a previously reported mass spectrometry-defined *NORAD* interactome comprised of 352 protein interactors^72^ and assessed their expression and dysregulation across iPSC-derived neurons, astrocytes, and microglia harboring the *MAPT* IVS10+16 mutation (**Figure 3A**). Comparable numbers of *NORAD* interactors were expressed in neurons (n=137), astrocytes (n=135), and microglia (n=138; **Figure 3A**; **Supplemental Table 21**). Notably, a substantial proportion of these *NORAD* interactors were differentially expressed in *MAPT* IVS10+16 cells, including 44 in neurons, 72 in astrocytes, and 67 in microglia (FDR≤0.05; **Figure 3B; Supplemental Figure 2A; Supplemental Table 22**), indicating enrichment of the *MAPT* mutation-associated transcriptional changes within the *NORAD* interaction network (Fisher’s exact test: neurons, p=2.2×10^−16^; astrocytes, p=9.487×10^−8^; microglia, p=2.2×10^−8^). While many dysregulated *NORAD* interactors were cell-type specific, 11 were significantly dysregulated across all three cell types (*ACTB, AGO1, ANXA2, ARPC2, CKAP4, G3BP1, GRSF1, MOV10, RBBP4, RBMX, WDR1*; **Figure 3C; Supplemental Figure 2A; Supplemental Table 22**), suggesting convergence on shared downstream pathways. Interestingly, *G3BP1*, *AGO1*, *MOV10*, and *ANXA2* are involved in stress granule formation, which has been implicated in FTLD-tau and other neurodegenerative diseases^73–76^. To further infer the functional significance of these interactions, we performed pathway enrichment analyses on dysregulated *NORAD* interactors from each cell type. Across neurons, astrocytes, and microglia, *NORAD*-associated proteins converged on pathways related to RNA biology, including the spliceosome and RNA degradation (**Supplemental Figure 2B–D**), consistent with established roles for *NORAD* in RNA regulatory processes^72^. Strikingly, the 11 shared dysregulated *NORAD* interactors were enriched for regulation of the actin cytoskeleton and stress granule assembly (**Supplemental Figure 2E**), implicating cytoskeletal organization and proteostasis as a potentially conserved downstream process associated with *NORAD* dysregulation across cell types^77, 78^.

**Figure 3:**
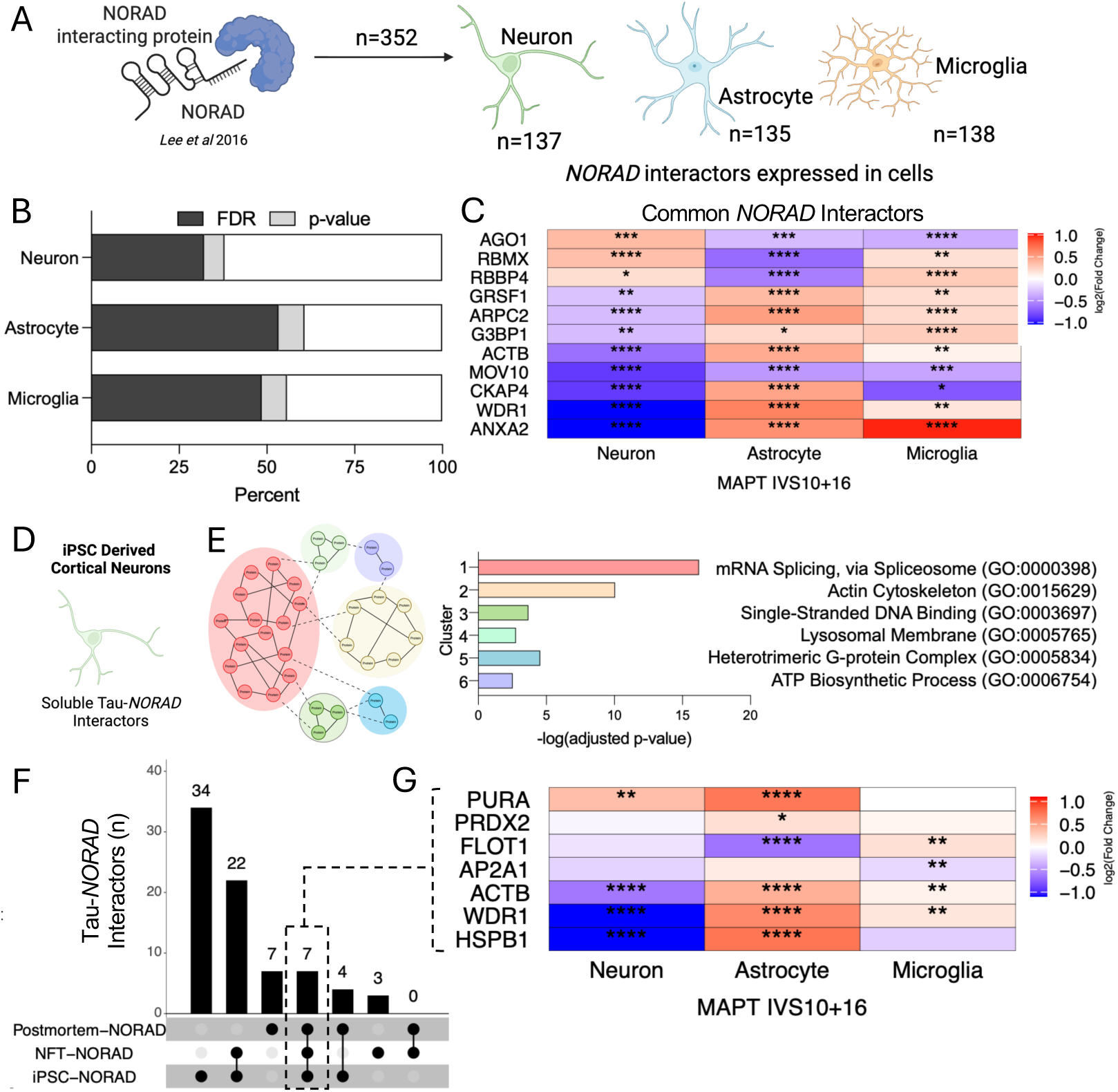
*NORAD*-associated protein networks converge on tau interactors and cytoskeletal regulatory pathways. A. *NORAD*-protein interactors were identified in Lee et al^72^ (n=352). A subset of NORAD-protein interactors were present at the mRNA level in iPSC-derived neurons (n=137), astrocytes (n=135), and microglia (n=138). B. *NORAD* interactors are differentially expressed in *MAPT* IVS10+16 neurons, astrocytes, and microglia. Dark gray, FDR≤0.05; light gray, p≤0.05; white, not significant. C. Heatmap of *NORAD* interactors that are differentially expressed across neurons, astrocytes, and microglia (FDR≤0.05). D. Schematic. Tau protein interactors were identified in iPSC-derived neurons (Kavanagh *et al*^79^) and were compared to *NORAD* interactors, revealing 67 shared interacting proteins. E. Clustering of 67 *NORAD*-tau interactors in iPSC-derived neurons was performed using the STRING Markov Cluster Algorithm (MCL) partitioned the network into six clusters. Clusters were then analyzed using KEGG pathway analysis to reveal significant enrichment of pathways related to RNA splicing and actin cytoskeleton. F. UpSet plot representing comparison of *NORAD*-protein interactors and Tau-protein interactors in iPSC neurons, postmortem brains, and neurofibrillary tangles reveals 7 shared protein interactors. G. Heatmap representing log_2_(fold-change) for RNA levels of the 7 shared *NORAD*- and tau-protein interactors in *MAPT* IVS10+16 neurons, astrocytes, microglia, and human brains. *, p≤0.05; **, p <0.01; ***, p< 0.001; ****, p< 0.0001.

Given the enrichment of genes dysregulated by *MAPT* IVS10+16 among *NORAD* interactors, we next asked whether *NORAD*-associated proteins intersect with tau-centered protein networks. Although tau itself was not identified as a direct *NORAD* interactor, multiple tau-associated proteins overlapped with the *NORAD* interactome. To define these relationships across distinct biochemical states of tau, we compared the *NORAD* interactome with previously reported tau interactors identified from three complementary datasets^79^: soluble tau interactors identified in iPSC-derived cortical neurons (n=1,648)^80^, soluble tau interactors identified from postmortem human brain tissue (n=698)^32, 81–83^, and insoluble tau interactors isolated from neurofibrillary tangles in human brain tissue (n=619)^83, 84^. The strongest overlap was observed with soluble tau interactors identified in iPSC-derived cortical neurons (n = 67; **Figure 3D–E**; **Supplemental Figure 3**; **Supplemental Table 23**), suggesting that *NORAD*-associated pathways intersect with soluble forms of neuronal tau. *NORAD* also shared interactors with soluble tau (n = 18; **Supplemental Figure 4**; **Supplemental Table 23**) and insoluble tau associated with neurofibrillary tangles (n = 32; **Supplemental Figure 5**; **Supplemental Table 23**) isolated from postmortem human brain tissue, suggesting that *NORAD*-related pathways extend beyond early soluble tau states and are related to disease pathology. These findings suggest that *NORAD*-associated protein networks are linked to both early soluble tau and end-stage aggregation-associated pathways, consistent with a persistent involvement across multiple stages of tauopathy.

To further characterize these overlapping networks, shared *NORAD*-tau interactors were clustered using the STRING Markov Cluster Algorithm (**Supplemental Figures 3-5**) followed by pathway enrichment analysis (**Figure 3E; Supplemental Figure 4B; Supplemental Figure 5B**). Interactors shared between *NORAD* and tau displayed cell-type specific and shared dysregulation across cell types (p≤0.05; **Supplemental Figure 3B-C; Supplemental Figure 4D-E; Supplemental Figure 5D-E**) suggesting common proteins or pathways with various tau interactors across neurons, astrocytes and microglia. Across all datasets, *NORAD*-tau interactors consistently converged on pathways related to the actin cytoskeleton, further implicating cytoskeletal regulation as a recurrent feature of *NORAD*-associated biology in tauopathy. Notably, seven genes were shared across *NORAD* and tau interaction networks in all datasets: *ACTB*, *AP2A1*, *FLOT1*, *HSPB1*, *PRDX2*, *PURA*, and *WDR1* (**Figure 3F**). Interestingly, these core interactors were also transcriptionally altered in *MAPT* IVS10+16 cells (**Figure 3G**). Among these, the cytoskeleton-associated genes, *ACTB* and *WDR1*, were significantly reduced in neurons and elevated in astrocytes and microglia (**Figure 3G**), mirroring the cell-type specific expression pattern observed for *NORAD* itself. Additionally, *HSPB1*, encoding Hsp27, a heat shock protein that prevents tau aggregation^85^, interacts with *NORAD*, and is significantly downregulated in neurons and upregulated in astrocytes (**Figure 3G**). Collectively, these findings place *NORAD* within tau-associated protein interaction networks linked to cytoskeletal organization and proteostasis across both soluble and insoluble tau states.

### MAPT IVS10+16 disrupts the NORAD–pumilio regulatory axis across CNS cell types

*NORAD* is a highly abundant and evolutionarily conserved lncRNA known for its role in regulating pumilio (PUM) proteins through direct molecular sequestration^19, 72, 86^. Under physiologic conditions, *NORAD* restrains pumilio activity by binding PUM1 and PUM2, limiting their interactions with pumilio response elements (PREs; 5′-UGUAHAUA) present within target mRNAs (**Figure 4A**)^72^. Pumilio proteins regulate mRNA stability and translation and have established roles in neuronal development and cellular homeostasis^19, 72, 87, 88^. Intriguingly, pumilio-regulated pathways intersect with biological processes increasingly implicated in neurodegeneration, including proteostasis, vesicular trafficking, autophagy, and cytoskeletal organization, raising the possibility that dysregulation of the *NORAD*-pumilio network may influence tau pathobiology. The dysregulation of *NORAD* across *MAPT* IVS10+16 cell types and human brain tissue led us to ask whether *PUM1* and *PUM2* mRNA expression were altered as a function of *MAPT* mutation. We discovered that both *PUM1* and *PUM2* were significantly increased in *MAPT* IVS10+16 neurons (*PUM1* p=0.001; *PUM2* p=3.66×10^−11^; **Figure 4B**). Conversely, *PUM1* was significantly reduced in astrocytes (p=0.0043) and *PUM2* was significantly reduced in microglia (p=0.018) (**Figure 4B**). Together, these finding point to cell-type specific alterations in the *NORAD*–pumilio regulatory axis in *MAPT* IVS10+16.

**Figure 4:**
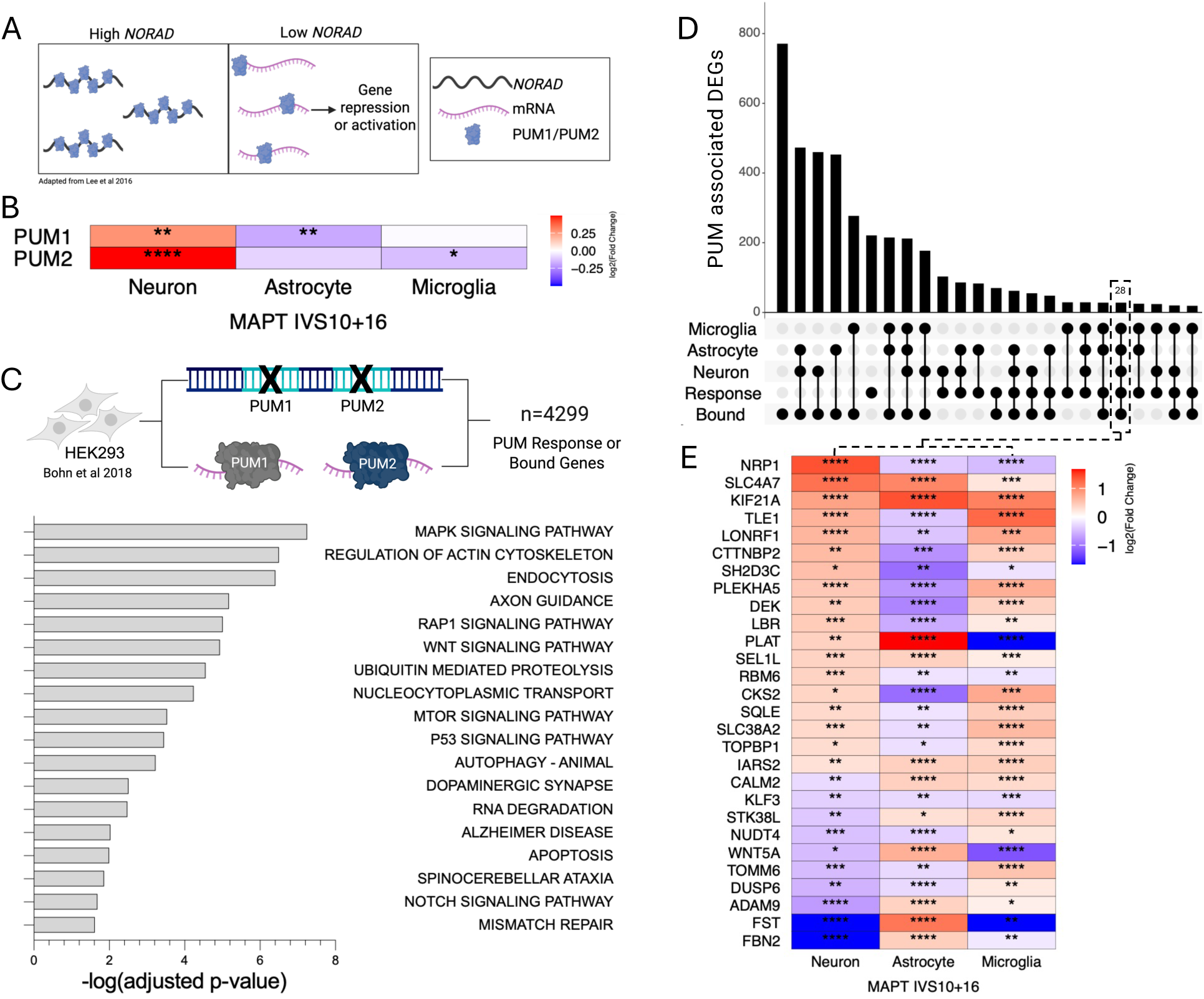
Pumilio proteins are dysregulated in *MAPT* IVS10+16 neurons, astrocytes, and microglia. A. Schematic of known *NORAD* and pumilio protein interaction adapted from^72^. High *NORAD* expression levels decrease pumilio target mRNA degradation/stabilizing function by binding to and sequestering pumilio. Low *NORAD* expression levels lead to hyperactive PUMILIO target mRNA degradation/stabilizing activity. B. Heatmap representing log_2_(fold-change) of *PUM1* and *PUM2* in *MAPT* IVS10+16 neurons, astrocytes, and microglia. *, p ≤ 0.05; **; p <0.01; ****, p < 0.0001. C. PUMILIO response (indirect) and bound (direct) genes were identified from Bohn *et al*^19^, revealing 4299 pumilio target genes. KEGG analysis was performed to reveal neurodegenerative related pathways, including regulation of actin cytoskeleton, ubiquitin mediated proteolysis, autophagy, Alzheimer’s disease, and others (adjusted p-value ≤ 0.05). D. Comparison of dysregulated pumilio response and bound genes in *MAPT* IVS10+16 neurons, astrocytes, and microglia (FDR ≤ 0.05 used as cutoff for differentially expressed genes). E. Heatmap representing log_2_(fold-change) of the 28 commonly dysregulated pumilio response and bound genes in *MAPT* IVS10+16 neurons, astrocytes, and microglia. *, p ≤ 0.05; **, p <0.01, ***, p < 0.001; ****, p < 0.0001.

To further define the potential downstream consequences of altered pumilio signaling in *MAPT* IVS10+16, we next examined genes regulated by pumilio^19^. We used datasets capturing both direct and indirect regulatory effects of pumilio proteins: direct pumilio targets (e.g. PUM-bound transcripts) were defined experimentally using RIP-Chip and PAR-CLIP approaches, while pumilio response genes were identified following combined *PUM1*/*PUM2* depletion and RNA-sequencing (e.g. transcripts regulated directly and indirectly by *PUM1* and *PUM2*)^19^. Together, these datasets defined 4,299 pumilio-associated genes^19^. Pumilio-associated genes revealed an enrichment in pathways associated with neurodegenerative disease, including AD, regulation of the actin cytoskeleton, endocytosis, ubiquitin-mediated proteolysis, and autophagy (**Figure 4C**). These findings suggest that pumilio-regulated networks converge on pathways broadly implicated in cytoskeletal organization and proteostatic regulation.

We next asked whether pumilio-regulated genes were altered in *MAPT* IVS10+16 neurons, astrocytes, and microglia. Differential expression analysis revealed widespread dysregulation of pumilio-associated transcripts across all three cell types (**Figure 4D; Supplemental Table 24**). Notably, 28 pumilio-associated genes were commonly dysregulated across neurons, astrocytes, and microglia (**Figure 4E**), defining a set of high-confidence pumilio regulatory targets altered in response to the *MAPT* IVS10+16 mutation.

Because *NORAD* canonically regulates pumilio activity (**Figure 5A**), we sought to further interrogate relationships within the *NORAD*-pumilio axis using targeted siRNA-mediated perturbation followed by quantitative PCR analysis in HEK293T cells. *NORAD* knockdown led to a modest increase in *PUM1* and a significant increase in *PUM2* (**Figure 5B**), suggesting that loss of *NORAD* may influence both pumilio activity and transcript abundance. Although *NORAD* is best known as a post-transcriptional regulator of pumilio proteins through molecular sequestration, these findings raise the possibility of feedback regulation within the *NORAD*-pumilio network. We next asked whether perturbation of pumilio proteins altered expression of other components within this regulatory network. Knockdown of *PUM1* resulted in a compensatory increase in *PUM2* expression, while *NORAD* transcript levels were not significantly altered (**Figure 5C**). Conversely, *PUM2* depletion reduced expression of both *PUM1* and *NORAD* (**Figure 5D**), suggesting coordinated regulation within the *NORAD*-pumilio axis. Considering the overlapping functions of pumilio proteins, we depleted both *PUM1* and *PUM2* simultaneously. Combined depletion of both *PUM1* and *PUM2* significantly reduced *NORAD* expression, suggesting reciprocal regulation between *NORAD* and pumilio proteins (**Figure 5E**). Together, these findings support coordinated dysregulation of the *NORAD*–pumilio regulatory axis in the setting of the *MAPT* IVS10+16 mutation (**Figure 5F**).

**Figure 5:**
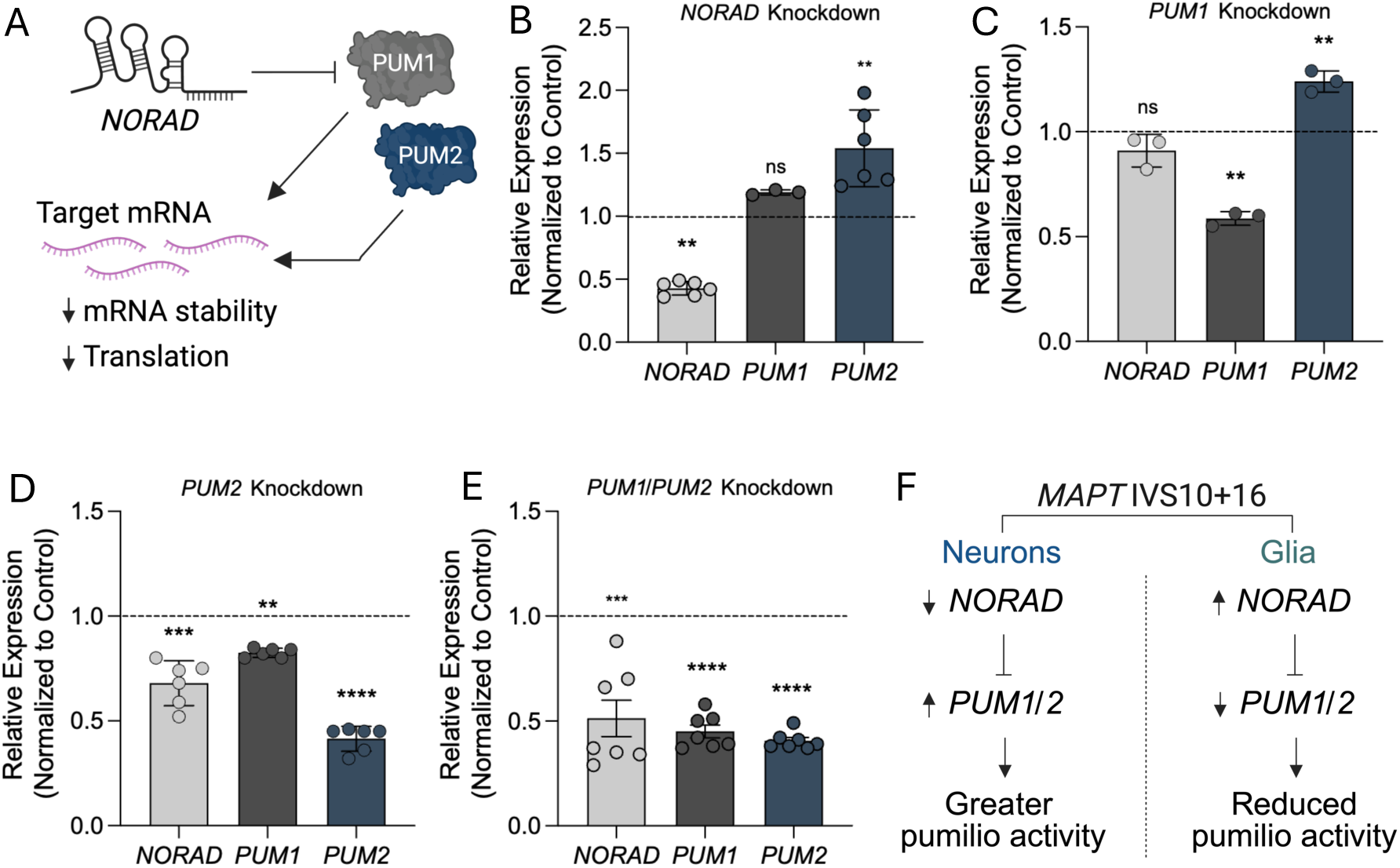
Perturbation of the *NORAD*–pumilio axis reveals reciprocal regulatory relationships. A. Schematic of the interaction between *NORAD* and Pumilio proteins. B. The effect of 72 h *NORAD* silencing in HEK293T cells. qPCR for *NORAD*, *PUM1,* and *PUM2* in siNORAD treated cells. C. The effect of 72 h *PUM1* silencing in HEK293T cells. qPCR for *NORAD*, *PUM1,* and *PUM2* in siPUM1 treated cells. D. The effect of 72 h *PUM2* silencing in HEK293T cells. qPCR for *NORAD*, *PUM1,* and *PUM2* in siPUM2 treated cells. E. The effect of 72 h *PUM1* and *PUM2* silencing in HEK293T cells. qPCR for *NORAD*, *PUM1,* and *PUM2* in siPUM1 and siPUM2 treated cells. F. Diagram of relationship between *NORAD* and *PUM1/2* in *MAPT* IVS10+16 neurons and glia. Data represent at least 3 biological replicates. Statistical analyses were conducted using Welch’s t-test; *, p≤0.05; **, p<0.001; ****, p<0.0001. Graphs represent mean ± SEM.

### The NORAD–pumilio regulatory axis modulates tau seeding susceptibility

Tau seeding is thought to occur when pathogenic tau templates the misfolding and aggregation of intracellular tau, a process believed to contribute to the progressive spread of tau pathology throughout the brain. Given the emerging relationship between the *NORAD*-pumilio axis and pathways implicated in tau propagation, we asked whether perturbation of this network influences susceptibility to tau aggregation. To test this, we utilized a HEK293T tau biosensor cell line stably expressing the repeat domain of tau containing the FTD-associated P301S mutation fused to CFP or YFP reporters^89^. Cells were treated with siRNA targeting candidate genes prior to exposure to seed-competent tau, followed by FRET-based quantification of tau aggregation (**Figure 6A**). Because *NORAD* emerged as a disease associated lncRNA dysregulated across *MAPT* IVS10+16 cell types and patient brain tissue, we first assessed the effects of *NORAD* depletion on tau seeding. *NORAD* knockdown significantly reduced tau seeding activity (**Figure 6B-C**), indicating that perturbation of *NORAD* is sufficient to alter susceptibility to tau aggregation. We next asked whether disruption of pumilio proteins similarly altered tau seeding phenotypes. Knockdown of either *PUM1* or *PUM2* individually resulted in significantly increased tau seeding (**Figure 6D-G**). To account for the partial redundancy of pumilio proteins, we depleted both *PUM1* and *PUM2* simultaneously and observed a similar significant increase in tau seeding activity (**Figure 6H-I**), consistent with a protective role for pumilio-associated pathways in limiting tau aggregation. Together, these findings identify the *NORAD*-pumilio regulatory axis as a modulator of tau seeding susceptibility and support a role for lncRNA-mediated RNA regulatory pathways in tau aggregation-associated phenotypes (**Figure 6J**). Interestingly, the effects of acute *NORAD* and pumilio perturbation on tau seeding differed from the expression patterns observed in *MAPT* mutant neurons and patient brain tissue, suggesting that the relationship between disease-associated *NORAD*-pumilio dysregulation and tau propagation may be biologically complex and potentially context dependent.

**Figure 6:**
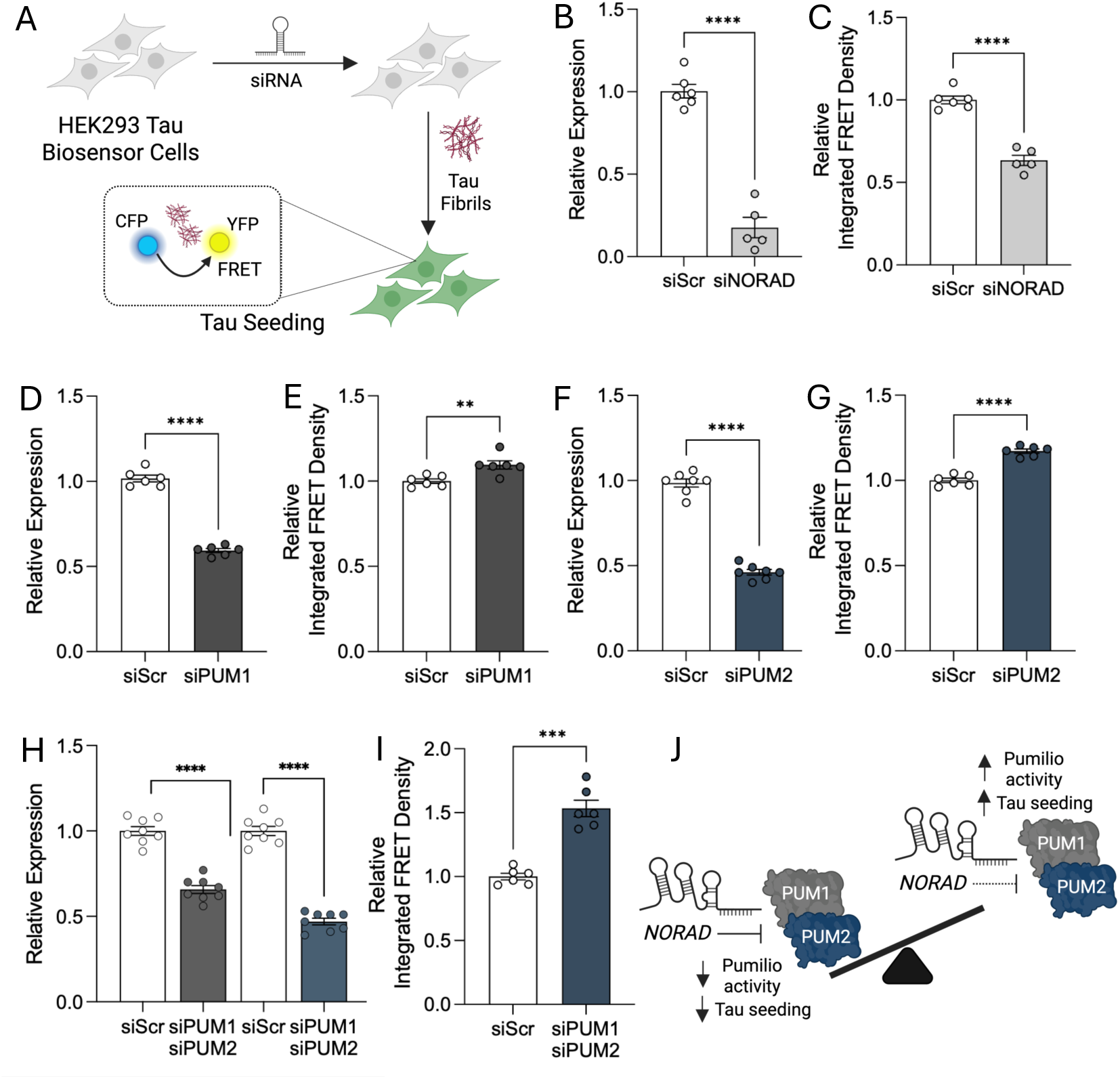
*PUM1*, *PUM2*, and *NORAD* regulate tau seeding in a tau biosensor cell line. A. Tau seeding workflow. Tau biosensor cells were treated with target gene siRNA for 18 hours followed by a media change. Seed competent tau were added using lipofectamine. Cells were collected after 24 hours, fixed using paraformaldehyde, and measured for FRET intensity using flow cytometry. B-C. Evaluating the impact of *NORAD* silencing on tau seeding in HEK293T biosensor model. B. qPCR for *NORAD* in siNORAD treated cells. C. Relative integrated FRET density (FRET MFI x percent FRET positive cells) normalized to vector control in siNORAD treated cells. D-E. Evaluating the impact of *PUM1* silencing on tau seeding in HEK293T biosensor model. D. qPCR for *PUM1* in siPUM1 treated cells. E. Relative integrated FRET density normalized to vector control in siPUM1 treated cells. F-G: Evaluating the impact of *PUM2* silencing on tau seeding in HEK293T biosensor model. F. qPCR for *PUM2* in siPUM2 treated cells. G. Relative integrated FRET density normalized to vector control in siPUM2 treated cells. H-I: Evaluating the impact of *PUM1* and *PUM2* silencing on tau seeding in HEK293T biosensor model. H. qPCR for *PUM1* and *PUM2* in siPUM1 and siPUM2 treated cells. I. Relative integrated FRET density normalized to vector control in siPUM1/siPUM2 treated cells. J. Summary of the effects of *NORAD* and *PUM1/2* on tau seeding. Data representative of at least 6 biological replicates. Statistical analyses were conducted using Welch’s t-test; *, *p* ≤ 0.05; **, *p* < 0.001; ****, *p* < 0.0001. Graphs represent mean ± SEM.

### The NORAD–pumilio regulatory network influences tau uptake pathways

Tau uptake is a critical step in the propagation of tau pathology, enabling extracellular tau aggregates to enter neighboring cells and seed intracellular aggregation^10, 90^. Although neurons represent the primary site of tau accumulation, astrocytes and microglia also contribute to tau propagation through uptake, clearance, inflammatory signaling, and release of tau species into the extracellular environment^49, 50^. Increasing evidence suggests that these cell-type specific processes collectively shape the spread of tau pathology throughout the brain. Because the *NORAD*–pumilio axis modulated tau seeding activity, we asked whether this network also influences cellular uptake of pathogenic tau species. To test this, HEK293T cells were treated with siRNAs targeting candidate genes followed by exposure to fluorescently labeled tau fibrils (Tau PFF-488), and intracellular tau uptake was quantified by flow cytometry (**Figure 7A**). Similar to its effects on tau seeding, *NORAD* depletion reduced tau uptake (**Figure 7B-C**). Knockdown of either *PUM1* or *PUM2* individually increased tau uptake (**Figure 7D-G**), consistent with the increased tau seeding observed following pumilio depletion. In contrast to the effects observed following individual paralog depletion, combined knockdown of *PUM1* and *PUM2* did not significantly alter tau uptake (**Figure 7H-I**), suggesting that coordinated disruption of pumilio signaling may differentially affect pathways involved in aggregate internalization relative to perturbation of individual paralogs. Together, these findings demonstrate that perturbation of the *NORAD*-pumilio regulatory axis influences multiple stages of tau pathology, including both aggregate uptake and intracellular seeding (**Figure 7J**). Given the established roles of neurons, astrocytes, and microglia in tau uptake, clearance, and propagation, these findings further support a model in which lncRNA-mediated RNA regulatory pathways contribute to multicellular mechanisms underlying the spread of tau pathology.

**Figure 7:**
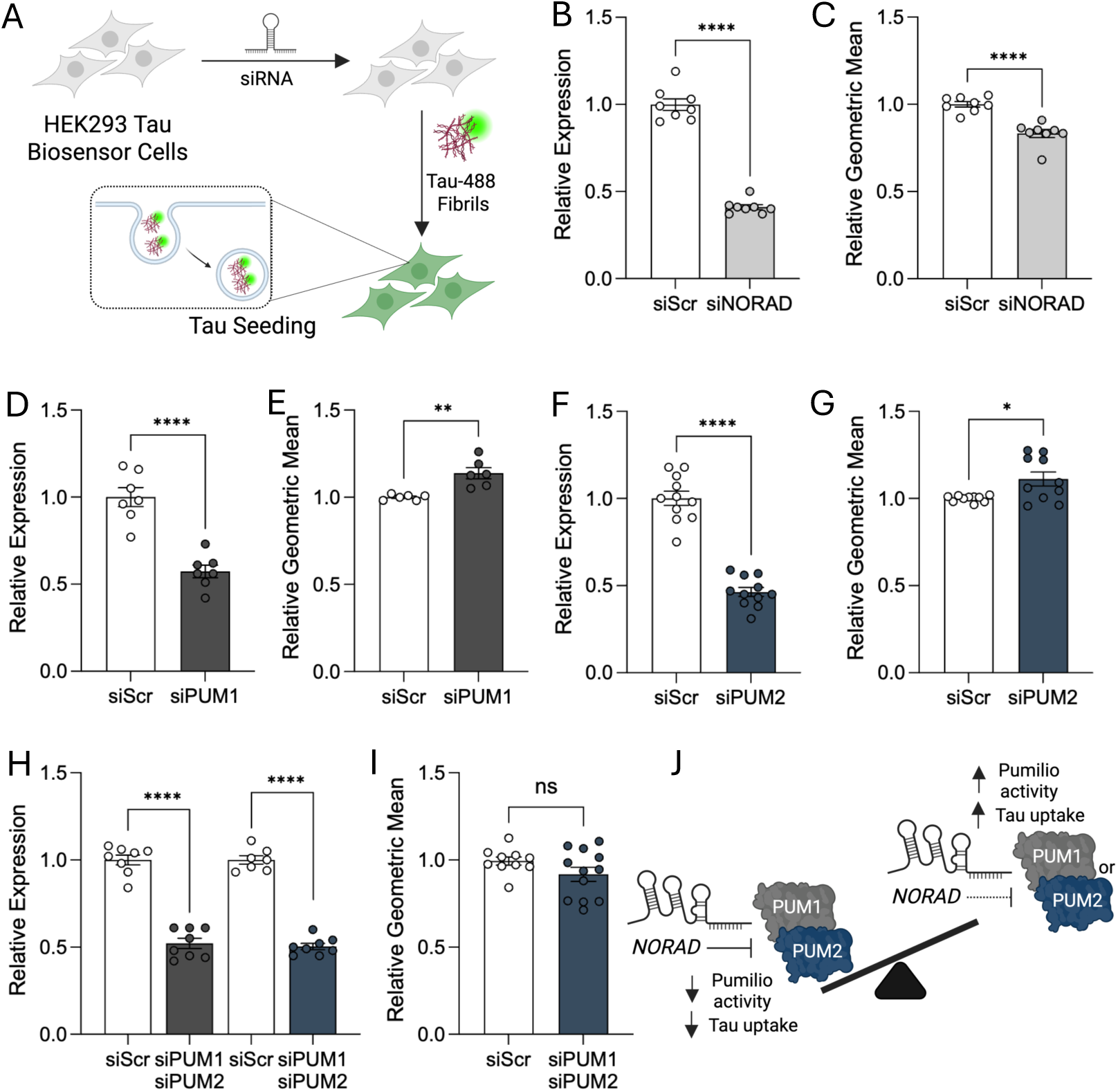
*PUM1*, *PUM2*, and *NORAD* regulate tau uptake *in vitro.* A. Tau uptake workflow. HEK293 cells were treated with target gene siRNA for 18 hours followed by a media change. Tau pre-formed fibrils conjugated to ATTO-488 were added to the media. Cells were collected after 24 hours and measured for 488+ signal using flow cytometry. B-C: Evaluating the impact of *NORAD* silencing on tau uptake in HEK293T cells. B. qPCR for *NORAD* in siNORAD treated cells. C. Relative geometric mean normalized to vector control in siNORAD treated cells. D-E: Evaluating the impact of *PUM1* silencing on tau uptake in HEK293T cells. D. qPCR for *PUM1* in siPUM1 treated cells. E. Relative geometric mean normalized to vector control in siPUM1 treated cells. F-G: Evaluating the impact of *PUM2* silencing on tau uptake in HEK293T cells. F. qPCR for *PUM2* in siPUM2 treated cells. G. Relative geometric mean normalized to vector control in siPUM2 treated cells. H-I: Evaluating the impact of *PUM1* and *PUM2* silencing on tau uptake in HEK293T cells. H. qPCR for *PUM1* and *PUM2* in siPUM1 and siPUM2 treated cells. I. Relative geometric mean normalized to vector control in siPUM1/siPUM2 treated cells. J. Summary of the effects of *NORAD* and *PUM1/2* on tau uptake. Data representative of at least 6 biological replicates. Statistical analyses were conducted using Welch’s t-test; *, *p* ≤ 0.05; **, *p* < 0.001; ****, *p* < 0.0001. Graphs represent mean ± SEM.

## Discussion

Long non-coding RNAs (lncRNAs) are increasingly recognized as important regulators of cellular identity and stress responses in the nervous system^55, 91, 92^, yet their roles in tauopathies remain incompletely understood. Here, we define the lncRNA landscape across iPSC-derived neurons, astrocytes, and microglia harboring the FTLD-associated *MAPT* IVS10+16 mutation and identify widespread cell type specific and shared lncRNA dysregulation. Among these altered lncRNAs, *NORAD* emerged as a priority candidate due to its broad dysregulation across neural cell types and patient brain tissue, enrichment in disease-relevant brain regions, and altered expression across multiple neurodegenerative diseases. Mechanistically, our findings position the *NORAD*-pumilio regulatory axis as a mediator of RNA homeostasis and tau propagation-associated phenotypes. Together, these data suggest that lncRNA-mediated regulatory pathways represent an important and underexplored layer of tauopathy biology.

A major finding of this study is the cell-type specificity of lncRNA expression and dysregulation in the context of a *MAPT* mutation. Although many lncRNAs were shared across neurons, astrocytes, and microglia, each cell type also expressed distinct lncRNA repertoires, with microglia exhibiting the largest set of cell-type restricted lncRNAs. This observation is consistent with prior studies showing that lncRNAs often display greater cell type specificity than protein-coding genes and contribute to specialized cellular states^14, 15, 93^. Notably, despite the expanded baseline lncRNA repertoire in microglia, neurons exhibited the greatest magnitude of *MAPT*-associated lncRNA remodeling. This pattern may reflect the central vulnerability of neurons to tau dysregulation, while also indicating that *MAPT* mutations alter broader RNA regulatory landscapes beyond protein homeostasis pathways alone. The enrichment of nuclear-localized lncRNAs across all cell types is further consistent with established roles for lncRNAs in chromatin organization, transcriptional regulation, and RNA processing^20, 94–97^.

Integration of iPSC-derived cell models with patient brain datasets identified *NORAD* and *MIR22HG* as the only lncRNAs commonly dysregulated across neurons, astrocytes, microglia, and *MAPT* IVS10+16 patient brain tissue. Among these, *NORAD* was prioritized based on its enrichment in brain regions vulnerable to FTLD-tau and its broader dysregulation across sporadic AD, autosomal dominant AD, and PD. *NORAD* has also been reported to be altered in biofluids from patients with FTD^98^, further supporting a relationship between *NORAD* dysregulation and neurodegenerative disease. Together, these findings suggest that *NORAD* dysregulation may extend beyond *MAPT*-associated FTLD-tau and represent a broader feature of neurodegenerative disease biology.

These findings extend prior work implicating lncRNAs in tauopathy and neurodegenerative disease^99–104^. Our previous studies showed that *MAPT* mutations broadly alter lncRNA expression in human iPSC-derived neurons and identified *SNHG8* as a regulator of stress granule biology through interactions with tau and RNA-binding proteins, including TIA1, FUS, and TDP43^21^. We also identified *FAM151B-DT* as a regulator of autophagy and proteostasis that modulates tau and α-synuclein aggregation through interactions with protein quality control pathways^22^. The present study extends this framework by defining lncRNA dysregulation across neurons, astrocytes, and microglia and by identifying *NORAD* as a disease-associated lncRNA that interfaces with pumilio-mediated RNA regulation, tau interaction networks, cytoskeletal pathways, and tau propagation-associated phenotypes^94, 99^.

*NORAD* is best characterized as a molecular decoy that restrains the activity of PUM1 and PUM2, two conserved RNA-binding proteins that regulate mRNA stability and translation^19, 72, 86, 105, 106–108^. In addition to this established sequestration mechanism, our perturbation studies suggest reciprocal regulatory interactions within the *NORAD*-pumilio axis. *NORAD* depletion increased *PUM* expression, whereas combined *PUM1/PUM2* depletion reduced *NORAD* levels, indicating that interactions between these molecules may extend beyond regulation of pumilio activity alone. Although the mechanisms underlying this feedback remain unresolved, these findings support a coordinated regulatory relationship between *NORAD* and pumilio proteins. Functionally, perturbation of this axis altered tau propagation-associated phenotypes: depletion of *PUM1* or *PUM2* increased tau seeding and tau uptake, whereas *NORAD* depletion reduced both processes. These opposing effects are consistent with the established antagonistic relationship between NORAD and pumilio proteins and suggest that pumilio-associated RNA regulatory pathways influence susceptibility to tau aggregation and propagation.

An important consideration is the relationship between disease-associated expression changes and acute perturbation phenotypes. In *MAPT* IVS10+16 neurons and patient brain tissue, *NORAD* expression was reduced while *PUM1* and *PUM2* expression were increased, a pattern broadly consistent with enhanced pumilio-associated signaling. This expression pattern is directionally consistent with our perturbation data, in which *NORAD* depletion increased *PUM* expression. However, the consequences of chronic dysregulation in human disease likely differ from those observed following acute perturbation in simplified cellular systems. Given the diverse roles of pumilio proteins in RNA homeostasis, translation, stress adaptation, and neuronal function, alterations in pumilio signaling may exert context-dependent effects during disease progression. The opposing expression patterns observed between neuronal and glial populations further suggest that the *NORAD*-pumilio axis may have distinct functions across neural cell types. These findings highlight the complexity of RNA regulatory pathways in neurodegenerative disease and suggest that *NORAD*-pumilio signaling may dynamically influence tau propagation-associated phenotypes across cellular and disease contexts.

Pumilio-regulated transcripts were enriched for pathways implicated in neurodegeneration, including autophagy, endocytosis, ubiquitin-mediated proteolysis, and regulation of the actin cytoskeleton^106^. These pathways are notable because tau propagation depends not only on intrinsic aggregation propensity, but also on cellular processes regulating vesicular trafficking, proteostasis, and cytoskeletal organization^33–35^. The observation that individual *PUM* depletion increased tau uptake whereas combined *PUM1/PUM2* depletion did not significantly alter uptake further supports a complex relationship between pumilio signaling and tau propagation. Given the partial redundancy and compensatory regulation between pumilio family members, simultaneous depletion of both proteins may engage distinct adaptive responses relative to perturbation of either paralog alone. Future studies will be required to define how specific pumilio-regulated RNA networks influence tau uptake and aggregation susceptibility.

Although the role of *NORAD* in neurodegeneration remains incompletely understood, emerging studies support its involvement in pathways relevant to neuronal homeostasis and disease. Loss of *Norad* in mice results in premature aging phenotypes, mitochondrial dysfunction, and multisystem degeneration associated with pumilio hyperactivity^59^. In neurodegenerative disease models, *NORAD* has been implicated in cellular stress responses in both PD and AD paradigms, including regulation of MPP+-induced neuronal injury^107^ and modulation of neprilysin/MME expression in Aβ-treated cells^108^. In prior work from our group, we also observed increased *NORAD* expression in Tau-P301L mice at 8 months of age, coinciding with the emergence of tau pathology^21^. Our findings extend this literature by linking *NORAD*-associated RNA regulation to tau uptake and seeding pathways relevant to tau propagation.

A complementary finding is the convergence of *NORAD*-associated protein networks with tau interaction networks across multiple biochemical states and disease stages. *NORAD* interactors showed the greatest overlap with soluble tau interactors identified in iPSC-derived neurons, suggesting that *NORAD*-associated pathways may be relevant to early neuronal tau biology before overt aggregation. Overlap with neurofibrillary tangle-associated proteins further suggests that *NORAD*-related pathways remain associated with later aggregated tau pathology. The smaller overlap observed in postmortem brain-derived soluble and insoluble tau interactors may reflect the biochemical and cellular heterogeneity of end-stage human tissue. Together, these findings suggest that *NORAD*-associated networks may remain engaged across multiple phases of tauopathy progression, spanning soluble tau dysregulation through aggregation-associated cellular remodeling.

In summary, we identify widespread lncRNA dysregulation across neurons, astrocytes, and microglia in the setting of *MAPT* mutation and nominate *NORAD* as a disease-associated lncRNA linked to neurodegenerative biology. This work highlights the importance of considering multicellular RNA regulatory mechanisms in models of tauopathy progression and identifies the *NORAD*-pumilio axis as a regulator of tau propagation-associated phenotypes. More broadly, these findings suggest that lncRNA-mediated regulatory networks contribute to the dynamic regulation of RNA homeostasis in tauopathy and may represent an important dimension of neurodegenerative disease progression.

## Experimental Procedures

### Patient consent

The informed consent was approved by the Washington University School of Medicine Institutional Review Board and Ethics Committee (IRB 201104178 and 201306108). The University of California San Francisco Institutional Review Board approved the operating protocols of the UCSF Neurodegenerative Disease Brain Bank (from which brain tissues were obtained). Participants or their surrogates provided consent for autopsy, in keeping with the guidelines put forth in the Declaration of Helsinki, by signing the hospital’s autopsy form. If the participant had not provided future consent before death, the DPOA or next of kin provided it after death. All data were analyzed anonymously.

### Induced Pluripotent Stem Cells (iPSC)

Human iPSCs used in this study were previously characterized^109^: *MAPT* IVS10+16 mutation carrier (GIH36C2) and CRISPR-Cas9 corrected isogenic control *MAPT* WT (GIH36C2d1D01). iPSCs cultured at 6% CO_2_ in mTeSR1 on Cultrex-coated tissue culture plates (R&D Systems, Cat. 3532-010-02) using Accutase™ (ThermoFisher Scientific, Cat. #NC9464543) for routine dissociation during passaging. Pluripotency of each line was confirmed by qPCR and immunocytochemistry (ICC)^109^. Chromosomal abnormalities were checked via karyotyping every 10 passages. Mutation status was confirmed by Sanger sequencing every 10 passages. iPSCs were maintained with less than 5% spontaneous differentiation and were free of mycoplasma contamination (tested monthly).

### Differentiation of iPSCs into cortical neurons and astrocytes

IPSCs were differentiated into neural progenitor cells (NPCs) as previously described (dx.doi.org/10.17504/protocols.io.x9cfr2w)^22, 56, 109, 110^. Briefly, neural aggregates were formed by plating 65,000 iPSCs per well in STEMdiff™ Neural Induction Medium (STEMCELL Technologies, Cat. #05835) in a 96-well v-bottom plate for 5 days. Cells were then transferred into 6 well plates. Neural rosettes were isolated using STEMdiff™ Neural Rosette Selection Reagent (STEMCELL Technologies, Cat. #05832) and cultured as neural progenitor cells (NPCs). NPCs were fed 2mL of STEMdiff™ Neural Induction Medium daily.

To generate cortical neurons, low passage NPCs were seeded at a density of 150,000 cells per well of a 12 well plate in STEMdiff™ Neural Induction Medium on Poly-L-Ornithine (Sigma, Cat. #P4957) and Laminin (Sigma, Cat. #L2020). After 24 hours, media was aspirated and changed to cortical differentiation medium: neurobasal medium (ThermoFisher, Cat. #21103049), B-27™ Supplement (Life Technologies, Cat. #17504-044), N-2 Supplement (1x; Thermofisher: 17502001), 20ng/mL BDNF (Sigma, Cat. #SR3014), 20ng/mL GDNF (Sigma, Cat. #G1777), 0.1mM cAMP (Sigma, Cat. #D0260), 1% Glutamax (ThermoFisher, Cat. #35050061), and 1% penicillin/streptomycin (ThermoFisher, Cat. #10378016). Media was changed every 2-3 days for 30 days.

To generate astrocytes, passage 3 NPCs were dissociated using 0.05% trypsin (Gibco, Cat. #25300-054) and replated in 6 well plates at a density of 60,000 cells/well in STEMdiff™ Neural Induction Medium. After 24 hours, media was aspirated and replaced with equal amounts of Astrocyte Medium (ScienCell, Cat. #1801). Media was changed every 2-3 days, and cells were split using 0.05% trypsin once cells reached 100% confluency. Cells were cultured as astrocytes for a total of 30 days then used for downstream experiments.

### Differentiation of iPSCs into microglia cells (iMGs)

To generate hematopoietic progenitor cells (HPCs), a STEMdiff™ Hematopoietic kit (STEMCELL Technologies, Cat. #05310) was used following manufacturer’s instructions as previously reported^57, 111–114^. Briefly, iPSCs were detached with ReLeSR™ (STEMCELL Technologies, Cat. #05872) and passaged in mTeSR1 supplemented with Rock inhibitor, Y-27632 Dihydrochloride (STEMCELL Technologies, Cat. #72302), to achieve a density of 45–80 aggregates/well. On day 0, cells were transferred to Medium A from the STEMdiff™ Hematopoietic Kit to pattern iPSCs towards mesoderm. On day 3, mesodermal cells were exposed to Medium B to further promote differentiation into HPCs, and cells remained in Medium B for 10 additional days. After 12 days in culture, HPCs were collected for fluorescence activated cell sorting (FACS) to enrich for CD43+ CD34+ HPCs (BioLegend, Cat. #343206, Clone: 10G7; BioLegend, Cat. #343504, Clone: 581).

Day 12 HPCs were plated onto Cultrex-coated 6 well plates in microglia media comprised of DMEM/F12 (Invitrogen, Cat. #11039021), 2X insulin-transferrin-selenite (Gibco, Cat. #41400-045), B-27™ Supplement (50X, 1: 25, Life Technologies, Cat. #17504-044), N-2 Supplement (100X, 1:200, ThermoFisher Scientific, Cat. #17502001), GlutaMAX (ThermoFisher, Cat. #35050061), 1X non-essential amino acids (Thermo Scientific, Cat. #11140050), 400 mM monothioglycerol (Sigma Aldrich, Cat. #M1753), 5 µg/mL human insulin (recombinant, Sigma Aldrich, Cat. #I2643), and freshly supplemented with maintenance cytokines (100 ng/mL IL-34 [PeproTech, Cat. #200-34], 50 ng/mL TGFβ1 [PeproTech, Cat. #100-21], and 25 ng/mL M-CSF [Peprotech, Cat. #300-25]). 1 million cells were grown on Cultrex-coated 6-well plates. Media was replenished every 2 days. On days 6 and 12, cells were split. Floating and adherent cells were evenly transferred from three wells into one new well. On day 25, cells were replated onto Cultrex-coated 6-well plates in microglia media supplemented with additional maturation cytokines (100 ng/mL CD200 [Novoprotein, Cat. #C311] and 100 ng/mL CX3CL1 [Peprotech, Cat. #300-31]). All iMGs used in this study were analyzed as mature iMGs harvested on day 28 post-HPC generation.

### Gene expression analysis in MAPT IVS10+16 iPSC-derived cells and patient brains

Bulk RNAseq was reanalyzed from iPSC-derived neurons, astrocytes, and microglia from *MAPT* IVS10+16 and isogenic controls using datasets described previously^21, 22, 56, 57^. Briefly, RNA was isolated using the RNeasy Mini Kit (Qiagen Cat. 74106) followed by ribodepletion (RiboZero). Samples were then sequenced by an Illumina HiSeq 4000 Systems Technology with a read length of 1 × 150 bp and an average library size of 36.5 ± 12.2 million reads per sample. Salmon (v. 0.11.3) was used to quantify the expression of the genes annotated within the human reference genome (GRCh38.p13)^115^. Protein coding transcripts and lncRNA transcripts were analyzed separately. Transcripts that were present in at least 10% of samples with expression of ≥ 0.1 TPM were included in subsequent analyses. Differential gene expression was performed using the DESeq2 (v.1.22.2) R package^116^.

Transcriptomic data from the middle temporal gyrus of patients with *MAPT* IVS10□+□16 (n=2) and neuropathology free controls (n=3) were previously described and reanalyzed in this work^21, 22, 56^. Briefly, RNA was isolated followed by ribodepletion. Samples were then sequenced by an Illumina HiSeq 4000 Systems Technology. Salmon (v. 0.11.3) was used to quantify the expression of the genes annotated within the human reference genome (GRCh38.p13)^115^. Protein coding transcripts and lncRNA transcripts were analyzed separately. Transcripts that were present in at least 10% of samples with expression of ≥ 0.1 TPM were included in subsequent analyses. Differential gene expression analysis was performed using the DESeq2 (v.1.22.2) R package^116^.

### RNA-sequencing in patient derived AD and PD brains

To assess *NORAD* and *MIR22HG* expression across neurodegenerative disease cohorts, publicly available bulk RNA-sequencing datasets from sporadic Alzheimer’s disease (sAD), autosomal dominant Alzheimer’s disease (ADAD), and Parkinson’s disease (PD) brains were analyzed (**Supplemental Table 25**)^117^. All cohorts included tissue from the frontal, parietal, and temporal cortices. The RNA library was prepared using a combination of either RiboZero Gold and TruSeq stranded RNA, or FastSelect ribosomal RNA blocking and FastSelect library prep.^117^ Sequencing was performed using one of Illumina HiSeq4000 or NovaSeq-6000 with 150 base pair paired-end reads. The data were aligned with STAR^118^ v 2.7.8a and counted with Salmon^115^ v 1.7.0. The tin.py script from RSeQC^119^ v 5.0.1 was run to calculate transcript integrity number (TIN) values for each transcript. FastQC v0.11.9 (*bioinformatics.babraham.ac.uk/projects/fastqc/*) and Picard v2.27.4 (*broadinstitute.github.io/picard/*) were implemented to produce quality control statistics, followed by summary of all analyses with multiqc^120^ v1.13.

A mixed effects model was run to analyze the difference in bulk transcript levels between cases and controls from multiple brain cortical regions (frontal, parietal, and temporal) using the lmer function from the program LmerTest^121^ v3.1.3.; the effect sizes were highly consistent with a parietal-only model for direction and magnitude of effect (r = 0.952, p = ≤ 6.000 × 10^−16^), indicating that merging multiple cortical regions into a single analysis did not substantially bias the results. Before analysis, transcriptomic principal component analysis (PCA) was run using prcomp (stats v4.4.0) to identify batch effects in the data. ComBat correction of raw counts based on sequencing batch was necessary for the brain cortex data using the sva^122^ v3.54.0 function ComBat_seq^123^. Counts were normalized before running DGA using the vst function from DESeq2 v1.46.0.^116^ Differential expression analyses for the ADAD cohort were performed using the following linear mixed effects model: normalized counts ∼ ADAD status + sex + postmortem interval + cortical region + median TIN + genetic principal components 1–2 + transcriptomic principal components 1–3 + (1 | individual ID). Similar models were used for the sAD and PD cohorts, with age at death additionally included as a covariate.

### Annotation of lncRNA genetic location

Genomic coordinates for lncRNAs were obtained using the Bioconductor package, biomaRt (v2.60.1), in RStudio (v4.4.1)^124, 125^. To account for annotation differences between genome builds, both the GRCh37 (Ensembl archive) and GRCh38 ( Ensembl v.114) assemblies were queried, and gene models were reconciled to ensure complete locus representation. For each lncRNA, chromosomal coordinates (chromosome, start position, end position, and strand) were extracted from the Ensembl hsapiens_gene_ensembl dataset. Ensembl gene identifiers were harmonized by removing version suffixes to ensure cross-assembly compatibility. To define cis-proximal genomic regions, lncRNA loci were extended by ±5 kb relative to annotated transcriptional boundaries. Protein-coding genes (Ensembl biotype = “protein_coding”) located within these extended regions were retrieved using the chromosomal_region filter in biomaRt. Genomic interval overlaps between extended lncRNA loci and protein-coding gene coordinates were identified using the GenomicRanges (v1.56.2) and IRanges (v2.38.1) packages in R, with overlaps computed via the findOverlaps function (type = “any”)^126^. For each overlapping pair, strand orientation and genomic boundaries were used to determine relative positional relationships and calculate base-pair distances between non-overlapping loci. lncRNAs were classified based on genomic co-localization with protein-coding genes within the ±5 kb window. lncRNAs were designated genic if they overlapped at least one protein-coding gene in the extended interval. lncRNAs were designated intergenic if no protein-coding genes were detected within the ±5 kb region flanking the lncRNA locus.

### LncRNA subcellular localization

LncRNA subcellular localization (cytoplasmic/nuclear) was determined using previously published data^60^. Cytoplasmic and nuclear RNA sequencing data from 15 different cell lines were extracted and analyzed for their cytoplasmic and nuclear enrichment^53^. Cytoplasmic-nuclear relative concentration index (RCI) was defined as: RCI= log_2_((Cytoplasmic expression (FPKM))/(Nuclear expression (FPKM))). RCI values for all lncRNAs were downloaded and imported into RStudio (v4.4.1)^125^. Common and cell type specific lncRNA RCI values were subsequently subset and used for analysis.

### LncRNA-protein interactions

To identify lncRNA-protein interactions, lncSEA 2.0 was used to determine experimentally validated and predicted protein interactions^61^. Protein interactions with an adjusted p-value ≤ 0.05 were used for downstream analyses. *NORAD* specific interactors were identified from Lee et al 2016^72^. Briefly, *NORAD*-interacting proteins were identified using previously published biotinylated RNA pull-down coupled with mass spectrometry datasets generated from conserved *NORAD* domains^72^.

### Gene ontology

Gene Ontology (GO) was determined using the Enrichr web server^127–129^. GO biological process, GO molecular function, GO cellular component, and KEGG analysis results were filtered (FDR ≤ 0.05) and manually selected for visual representation. - Log_2_(adjusted p-value) was used for data visualization.

### GTEx analysis

Bulk tissue RNA-sequencing results were accessed through The Genotype-Tissue Expression (GTEx) Project^130^. One file per tissue type was downloaded and subsequently loaded into and read by RStudio. *NORAD* and *MIR22HG* transcripts per million were subset by tissue type and exported as an Excel file using the writexl package^131^. The data used for the analyses described in this manuscript were obtained from the GTEx Portal on 09/25/2025 (V11).

### Total RNA isolation

RNA was isolated using the Qiagen RNeasy Mini Kit (Qiagen, Cat. #74106). Briefly, cell pellets were resuspended in QIAzol Lysis Reagent (Qiagen, Cat. #79306) and incubated at room temperature for 5 minutes. Chloroform was then added, followed by inversion until mixed, and incubated at room temperature for 2 minutes. Tubes were then centrifuged at 15,000xg for 15 minutes at 4°C. Tubes were then transferred to ice followed by careful extraction of the top aqueous layer of supernatant into a new tube. Equal amounts of 70% ethanol were added to the aqueous layer followed by vortexing for 15 seconds. Samples were then added to RNeasy Mini Spin Columns and centrifuged at 8000xg for 1 minute. Supernatant was discarded followed by the addition of Buffer RW1 to spin column. Samples were again spun at 8000xg for 1 minute. To remove genomic DNA, 80µL of RDD-diluted RNase Free DNase (Qiagen, Cat. #79254) was added to each column and incubated at room temperature for 15 minutes. Buffer RW1 was then added to spin column and centrifuged at 8000xg for 1 minute followed by discard of flow through. Buffer RPE was added to spin column followed by centrifugation at 8000xg for 1 minute and discard of flow through. This was repeated and followed by an additional centrifugation at 8000xg for 1 minute to dry resin. To elute RNA, spin columns were transferred to clean Eppendorf tubes, followed by addition of RNase free water directly to resin and incubated at room temperature for 5 minutes. Samples were centrifuged at 8000xg for 1 minute. To assess RNA purity and concentration, each sample was measured using the NanoDrop™ 8000 Spectrophotometer.

### Quantitative PCR (qPCR)

RNA expression was analyzed by real-time PCR. *NORAD*, *PUM1*, *PUM2*, and *GAPDH* were measured using iTaq™ Universal SYBR® Green Supermix (Bio-Rad, Cat. #172-5120). Primer sequences were as follows: *NORAD* 5’-AAGCTGCTCTCAACTCCACC-3’ and 5’-GGACGTATCGCTTCCAGAGG-3’; *PUM1* 5’-GCATTTGGACAAGGTCTGGCAG-3’ and 5’-GCTACAAGTCGAACAGGAGCTC-3’; *PUM2* 5’-CGGTTAATGGCTCCAACACCTG-3’ and 5’-CGAAACAGACCATTTGTGCTGCC-3’; *GAPDH* 5’-TGCACCACCAACTGCTTAGC-3’ and 5’-GGCATGGACTGTGGTCATGAG-3’. Samples were run with at least two technical replicates with *GAPDH* as a housekeeping gene for normalization. The QuantStudio™ 3 or 12K machine were used for qPCR. Analysis was conducted using the comparative CT method. Technical replicate CT values were averaged and normalized to *GAPDH*. Technical replicates with averages of a standard error of 20% or higher were re-run.

### Pumilio targets

Pumilio targets were sourced from Bohn *et al* 2018^19^. Briefly, pumilio targets were defined as bound and response genes. Pumilio bound targets were defined as those mRNAs bound to *PUM1* or *PUM2* in HeLA or HEK293 cells identified using RIP-ChIP and PAR-CLIP assays from previously reported databases^132–134^. Pumilio response genes were defined as those changed as a function of *PUM1*/*PUM2* double knockdown in HEK293 cells. Pumilio bound and response genes lists were compared to those protein coding dysregulated genes in *MAPT* IVS10+16 neurons, astrocytes and microglia. Genes with FDR≤0.05 were used in subsequent analyses.

### Tau interactor analysis

Tau interactors were defined in a meta-analysis from Kavanagh et al 2022^79^. Tau interactors identified in iPSC derived neurons (soluble tau), postmortem human AD brains (total n= 16; soluble tau) and healthy control brains (total n= 11; soluble tau), and NFTs isolated from human AD brains (total n=11; insoluble tau) were used for subsequent analyses^32, 80–84, 135^.

### Plasmids and transient transfection

To evaluate the impact of *NORAD*, *PUM1*, and *PUM2* silencing, dicer-substrate and scrambled siRNAs were synthesized (IDT). HEK293T or HEK293T-tau biosensor cells were transiently transfected with siRNAs (*NORAD, PUM1, or PUM2* single knockdowns: 10nM; *PUM1*/*PUM2* double knockdown: PUM1 10nM, PUM2 20nM).

Transfection was performed using Lipofectamine RNAiMAX (Invitrogen, Cat. #13778100) following manufacturer’s instructions. Briefly, 75,000-150,000 cells were seeded onto Poly-D-Lysine (ThermoFisher, Cat. #A3890401) coated 12 well plates. After 24 hours, siRNA was mixed with 2.5µL of Lipofectamine RNAiMAX in 750µL of Opti-MEM (Gibco, Cat. #31985062) and incubated at room temperature for 15 minutes. Cell media was then aspirated, and siRNA mixture was added to cells for 16 hours and cells were evaluated 48 hours post transfection.

### Tau uptake assay

HEK293 cells were cultured at 37°C Celsius at 5% CO_2_ in DMEM (Gibco, Cat. #11965-092) supplemented with 10% fetal bovine serum (Biowest, Cat. #S1620) and 1% penicillin-streptomycin (ThermoFisher, Cat. #15140-122). Cells were split when they reached ∼80% confluency by washing with DPBS (Gibco, Cat. #14190-144) and dissociated with 0.05% trypsin (Gibco, Cat. #25300-054). Briefly, 75,000 cells were seeded into each well of a 12-well plate. After 24 hours, cells were transfected with *NORAD*, *PUM1*, or *PUM2* siRNAs (10nM), overexpression plasmids (*NORAD* 250ng), control siRNA, or control plasmids in OptiMEM (Gibco, Cat. #31985062). Media was changed after 24 hours to DMEM supplemented with 10% fetal bovine serum and 1% penicillin-streptomycin. After 24 hours, 10µM of human recombinant Tau441 (2N4R) fibrils conjugated to ATTO-488 (StressMarq, Cat. #SPR-329-A488) was added to each well. After 24 hours, cells were harvested with 0.05% trypsin and prepared for flow cytometry by resuspending each cell pellet in DPBS supplemented with 1% fetal bovine serum and 0.2% 0.5M EDTA (BioLogical Molecular Reagents, Cat. #5109.500). Samples were analyzed on the LSR Fortessa™ flow cytometer. For each experiment, 10,000 cells per replicate were analyzed, with data processed using FlowJo v10 software.

### Tau seeding assay

HEK293T cells were cultured at 37°C at 5% CO_2_ in DMEM supplemented with 10% fetal bovine serum (Biowest, Cat. #S1620) and 1% penicillin-streptomycin (ThermoFisher, Cat. #15140-122). Cells were split when they reached ∼80% confluency by washing with DPBS and dissociated with 0.05% trypsin (Gibco, Cat. #25300-054). Tau seeding assays were conducted using a protocol previously described^89^. Briefly, this assay uses HEK293T cells expressing the tau repeat domain with the FTD-linked P301S variant of tau fused to either CFP or YFP^89^. Cells were seeded at a density of 75,000 cells per well of a 12 well plate. After 24 hours, cells were transfected with *NORAD*, *PUM1*, or *PUM2* siRNAs (10nM), overexpression plasmids (*NORAD* 250ng), control siRNA, or control plasmids in Opti-MEM (Gibco, Cat. #31985062). Media was changed after 24 hours to DMEM supplemented with 10% fetal bovine serum and 1% penicillin-streptomycin (ThermoFisher, Cat. #15140-122). After an additional 24 hours, 10µM of recombinant Tau-441 (2N4R) P301S Mutant Pre-formed Fibrils (StressMarq, Cat. #SPR-471) were transfected using 5uL of Lipofectamine 2000 (ThermoFisher, Cat. #11668019). After 24 hours, cells were dissociated with 0.05% trypsin and fixed using 4% paraformaldehyde (Electron Microscopy Services, Cat. #15710) for 10 minutes and resuspended in flow cytometry buffer (1% fetal bovine serum and 0.2% 0.5M EDTA [BioLogical Molecular Reagents, Cat. #5109.500]). Flow cytometry was conducted using the BD LSR Fortessa™ Cell Analyzer (BD Biosciences). Gating strategy was conducted as described in Banning et al^136^. For each experiment, 10,000 cells per replicate were analyzed, with data processed using FlowJo v.10 software.

### Statistical analyses

DESeq2 (v.1.22.2) was used to determine differential gene expression^116^. For statistical analyses between two groups, an unpaired Welch’s t-test was used to compare differences between two groups. Among three or more groups, a one-way ANOVA was used. Welch’s t-tests and one-way ANOVAs were determined using GraphPad PRISM software (v.10). Tukey HSD tests were used to perform pairwise comparisons where indicated. P-values≤0.05 were considered statistically significant.

## Acknowledgments

We thank the research subjects and their families who generously participated in this study. We thank Torri Ball for thoughtful discussions. This work was supported by access to equipment made possible by the Hope Center for Neurological Disorders, the Neurogenomics and Informatics Center, and the Departments of Neurology and Psychiatry at Washington University School of Medicine. Funding provided by the National Institutes of Health (P30 AG066444, RF1 NS110890, U54 NS123985), Rainwater Charitable Organization (CMK), Farrell Family Fund for Alzheimer’s Disease (CMK), and UL1TR002345. The Genotype-Tissue Expression (GTEx) Project was supported by the Common Fund of the Office of the Director of the National Institutes of Health, and by NCI, NHGRI, NHLBI, NIDA, NIMH, and NINDS. Diagrams were generated using BioRender.com.

Data collection and sharing for this project was supported by The Dominantly Inherited Alzheimer Network (DIAN, U19AG032438 & 3U19AG032438-12S2) funded by the National Institute on Aging (NIA), the Alzheimer’s Association (SG-20-690363-DIAN LATAM & DIAN OBS-26-1552363), the German Center for Neurodegenerative Diseases (DZNE), Raul Carrea Institute for Neurological Research (FLENI), Partial support by the Research and Development Grants for Dementia from Japan Agency for Medical Research and Development (AMED), the Korea Health Technology R&D Project through the Korea Health Industry Development Institute (KHIDI), Korea Dementia Research Center (KDRC), funded by the Ministry of Health & Welfare and Ministry of Science and ICT, Republic of Korea RS-2024-00344521, and Spanish Institute of Health Carlos III (ISCIII). This manuscript has been reviewed by DIAN Study investigators for scientific content and consistency of data interpretation with previous DIAN Study publications. We acknowledge the altruism of the participants and their families and contributions of the DIAN research and support staff at each of the participating sites for their contributions to this study.

## Data Availability

All data generated or analyzed during this study are included in this published article and its supplementary information files.

## Author Contributions

Designed experiments: JZ, CMK. Performed and analyzed experiments: JZ, GH, ES, MB, JM, AR, BP, TM, MM, CC, AIK, CMK. Provided experimental tools: DIAN. Provided funding: CMK. Wrote the manuscript: JZ, CMK. Revised and approved manuscript: JZ, GH, ES, MB, JM, AR, BP, TM, MM, CC, AIK, CMK.

