## Supplemental Figures for "The *NORAD*-pumilio regulatory axis links lncRNA dysregulation to tau propagation-associated phenotypes"

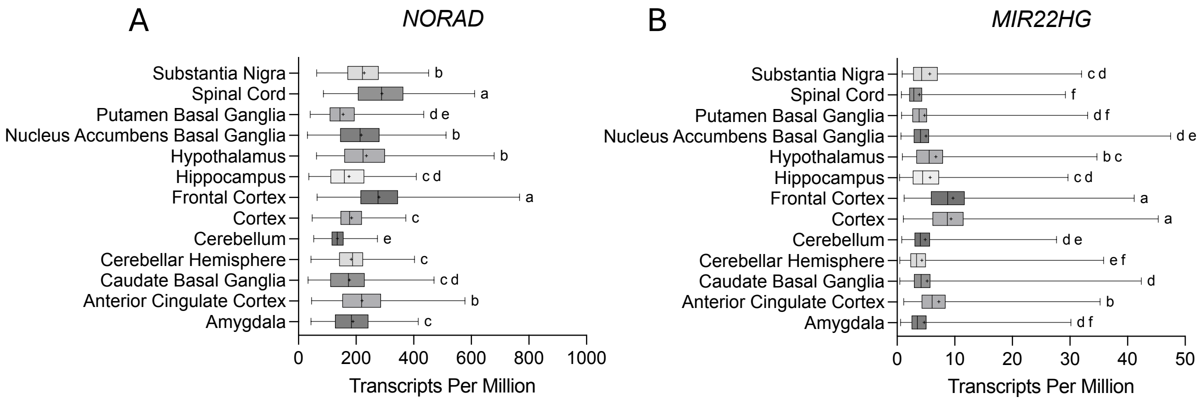


**Supplemental Figure 1: *NORAD* and *MIR22HG* expression across brain regions.** A. *NORAD* expression levels (transcripts per million) across brain regions indicate significant enrichment in the frontal cortex and spinal cord. Letters denote results from Tukey's HSD post hoc test; different letters indicate significant differences among means (FDR≤0.05). B. *MIR22HG* expression levels (transcripts per million) across brain regions indicate significant enrichment in the cortex. Letters denote results from Tukey's HSD post hoc test, indicating significant differences among means (FDR≤0.05). Data was obtained from GTEx V11.

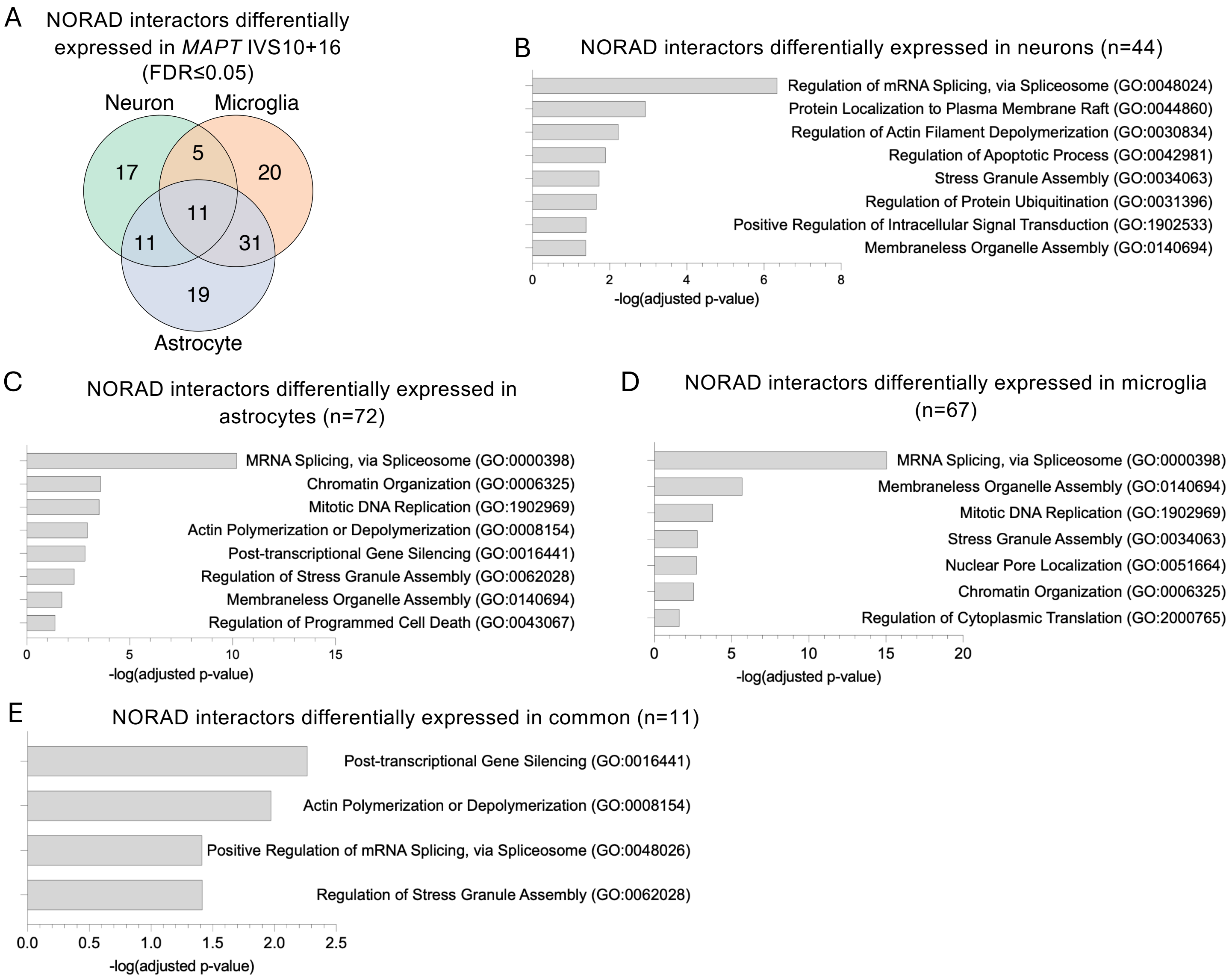


**Supplemental Figure 2: *NORAD* interactors are dysregulated in neurons, astrocytes, and microglia.** *NORAD*-protein interactors were identified in Lee *et al* 2016 [72](n=352). A. Venn diagram representing the number of cell type specific and shared dysregulated *NORAD*-protein interactors (FDR ≤ 0.05). B-E: Cell type-specific and shared KEGG pathway analysis. Plotted values represent -log(adj.p-value), providing a more intuitive visualization of KEGG pathway significance.

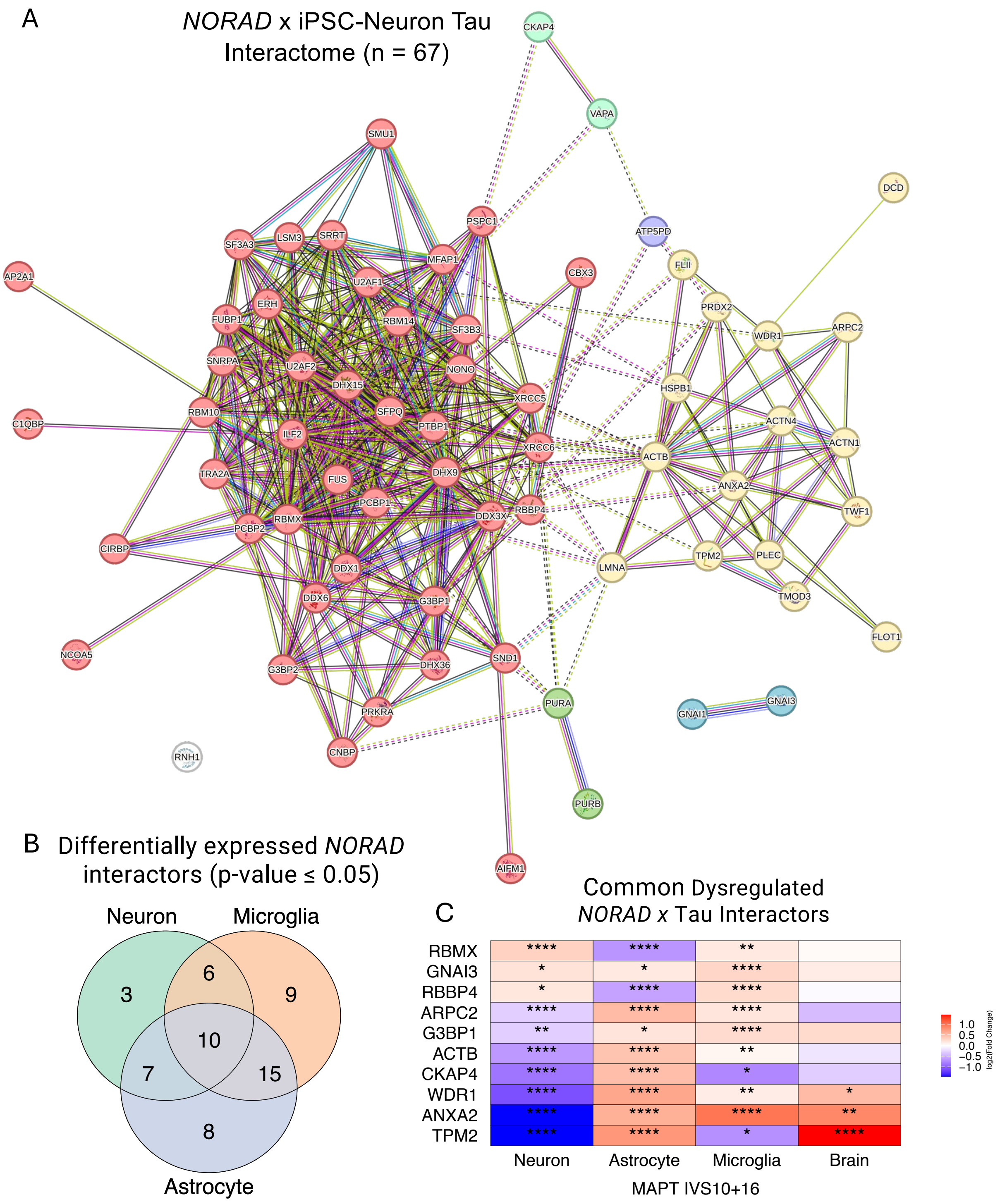


**Supplemental Figure 3: Shared *NORAD* and iPSC-derived neuron tau interactors are dysregulated in iPSC-derived neurons, astrocytes and microglia.** A: STRING network analysis of all *NORAD*-iPSC tau protein interactors (n=67). Solid lines indicate interaction between proteins. Dotted lines indicate interaction between edges of cluster. Clusters determined by Markov cluster algorithm. B: Venn diagram representing the number of cell type specific and shared dysregulated ($p \leq 0.05$) *NORAD*-protein interactors. C: Heatmap representing $\log_2$(fold-change) of 10 dysregulated ($p \leq 0.05$) *NORAD*-protein interactors in iPSC derived neurons, astrocytes and microglia; *, $p \leq 0.05$; **, $p < 0.001$; ****, $p < 0.0001$.

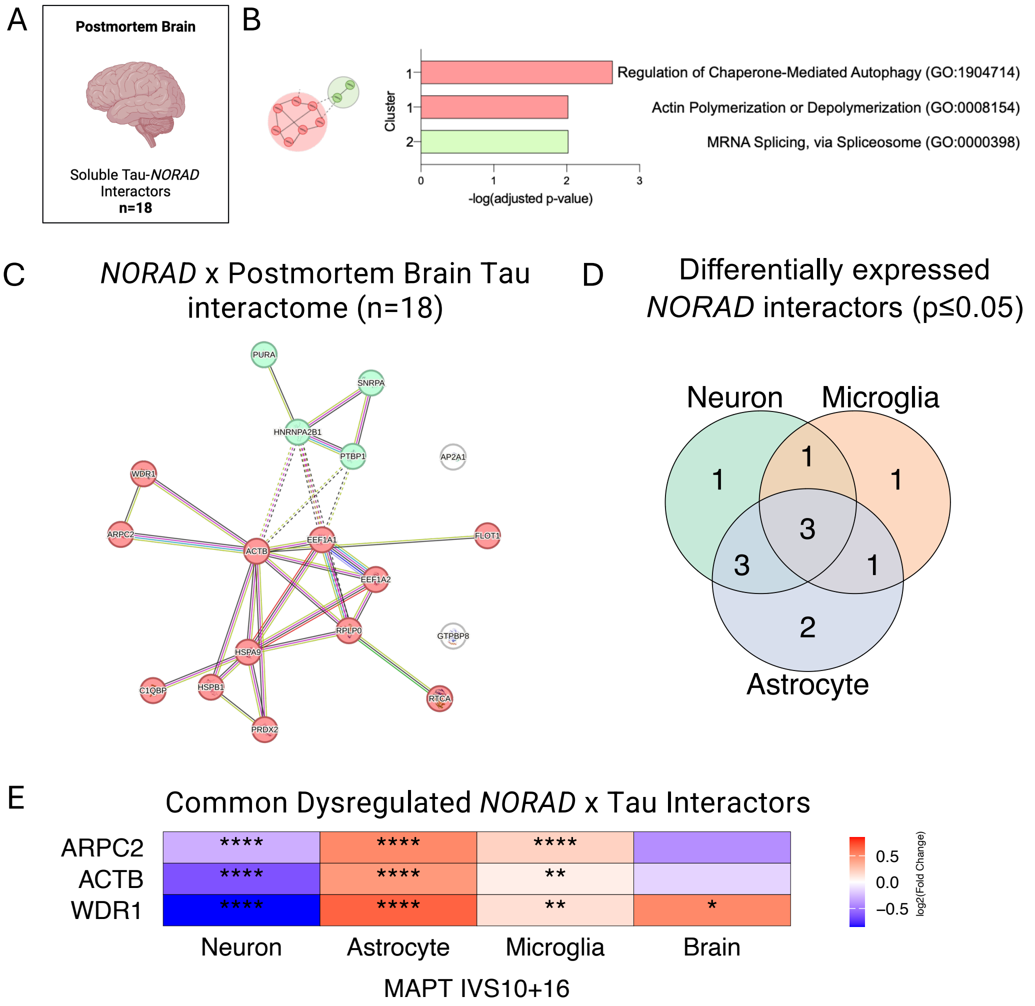


**Supplemental Figure 4: Shared *NORAD* and postmortem brain tau interactors are dysregulated in iPSC-derived neurons, astrocytes and microglia**. A: Tau protein interactors in the detergent soluble fraction of Alzheimer's disease brains revealed 18 protein interactors shared with *NORAD*. B: Clustering of 18 *NORAD*–tau interactors using the STRING MCL partitioned the network into three clusters. Clusters were then analyzed using KEGG pathway analysis to reveal significant enrichment of pathways related to actin cytoskeleton. C: STRING network analysis of all NORAD-postmortem tau protein interactors (n=18). Solid lines indicate interaction between proteins. Dotted lines indicate interaction between edges of cluster. Clusters determined by Markov cluster algorithm. D: Venn diagram representing the number of cell type specific and shared dysregulated *NORAD*-protein interactors ($p \leq 0.05$). E: Heatmap representing $\log_2$(fold-change) of 3 commonly dysregulated *NORAD*-protein interactors in iPSC derived neurons, astrocytes and microglia; *, $p \leq 0.05$; **, $p < 0.001$; ****, $p < 0.0001$.

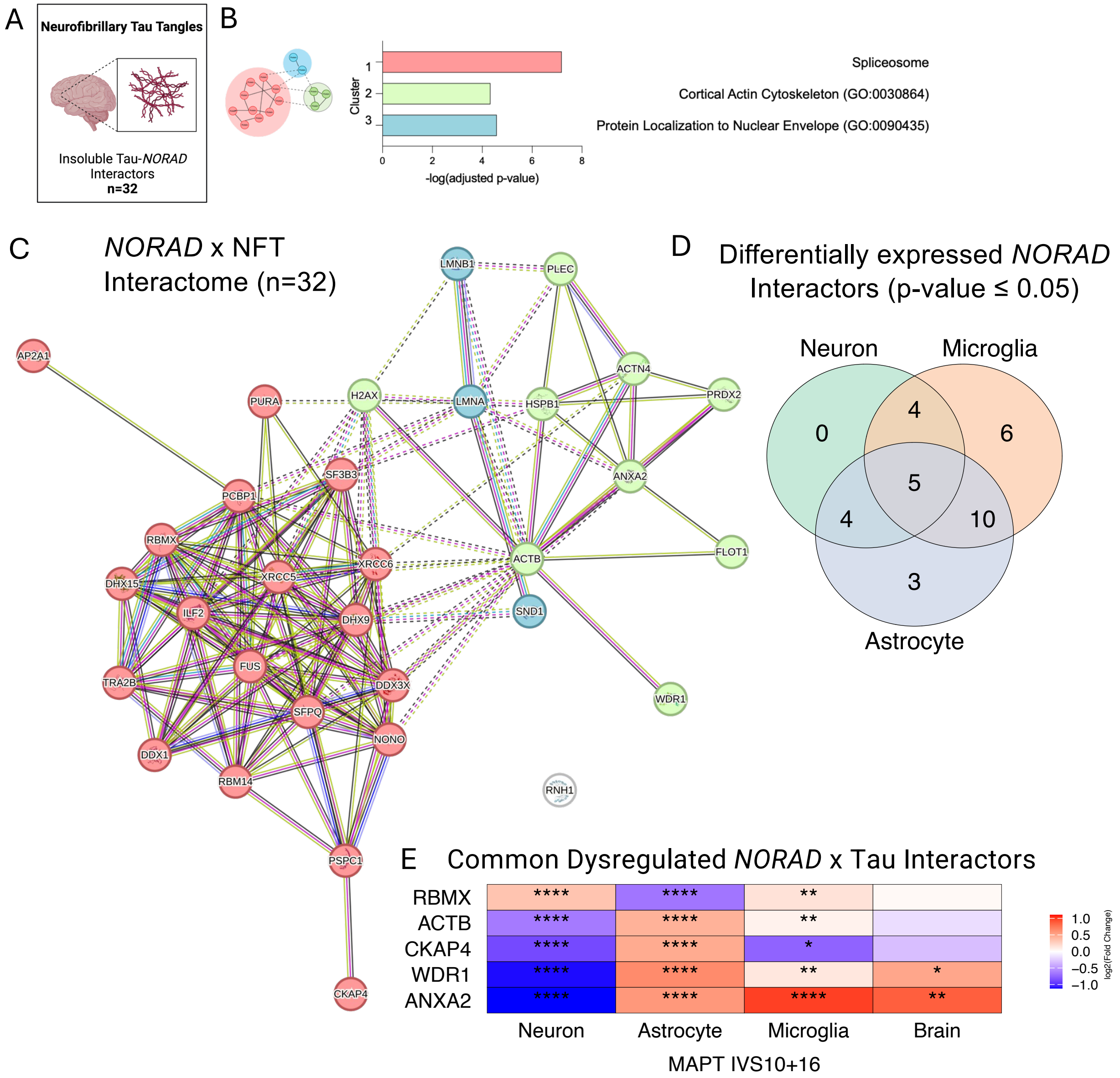


**Supplemental Figure 5: Shared *NORAD* and NFT interactors are dysregulated in iPSC-derived neurons, astrocytes and microglia.** A: Tau protein interactors were identified in neurofibrillary tau tangles isolated from Alzheimer's disease brains and were found to overlap with 32 *NORAD*-protein interactors. B: Clustering of 32 *NORAD*-tau interactors using the STRING MCL partitioned the network into three clusters. Clusters were then analyzed using KEGG pathway analysis to reveal significant enrichment of pathways related to the spliceosome and actin cytoskeleton. C: STRING network analysis of all *NORAD*-neurofibrillary tangle interactors (n=32). Solid lines indicate interaction between proteins. Dotted lines indicate interaction between edges of cluster. Clusters determined by Markov cluster algorithm. D: Venn diagram representing the number of cell type specific and shared dysregulated *NORAD*-protein interactors (p≤0.05). E: Heatmap representing $\log_2$(fold-change) of 5 commonly dysregulated *NORAD*-protein interactors in iPSC derived neurons, astrocytes and microglia; *, $p \leq 0.05$; **, $p < 0.001$; ****, $p < 0.0001$.
